# *ppe51* variants promote non-replicating *Mycobacterium tuberculosis* to grow at acidic pH by selectively promoting glycerol uptake

**DOI:** 10.1101/2021.05.19.444820

**Authors:** Shelby J. Dechow, Jacob J. Baker, Megan R. Murto, Robert B. Abramovitch

## Abstract

In defined media supplemented with single carbon sources, *Mycobacterium tuberculosis* (Mtb) exhibits carbon source specific growth restriction. When supplied glycerol as the sole carbon source at pH 5.7, Mtb establishes a metabolically active state of nonreplicating persistence known as acid growth arrest. We hypothesized that acidic growth arrest on glycerol is not a metabolic restriction, but rather an adaptive response. To test this hypothesis, we conducted forward genetic screens that identified several Mtb mutants that could grow under these restrictive conditions. All of the mutants were mapped to the *ppe51* gene and resulted in three amino acid substitution – S211R, E215K, and A228D. Expression of the PPE51 variants in Mtb promoted growth at acidic pH showing that the mutant alleles are sufficient to cause the dominant gain-of-function, enhanced acid growth (*eag*) phenotype. Testing growth on other single carbon sources showed the PPE51 variants specifically enhanced growth on glycerol, suggesting *ppe51* plays a role in glycerol uptake. Using radiolabeled glycerol, enhanced glycerol uptake was observed in Mtb expressing the PPE51 (S211R) variant, with glycerol overaccumulation in triacylglycerol. Notably, the *eag* phenotype is deleterious for growth in macrophages, where the mutants have selectively faster replication and reduced in virulence in activated macrophages as compared to resting macrophages. Recombinant PPE51 protein exhibited differential thermostability in the WT or S211R variants in the presence of glycerol, supporting the *eag* substitutions alter PPE51-glycerol interactions. Together, these findings support that PPE51 variants selectively promote glycerol uptake and that slowed growth at acidic pH is an important adaptive mechanism required for macrophage pathogenesis.

## Introduction

During infection*, Mycobacterium* tuberculosis (Mtb) senses and adapts to a variety of immune cues including hypoxia^1,2^, nutrient starvation^3,4^, pH changes^5^, and nitrosative and oxidative stress^6^. Exposure to these stresses can promote Mtb to establish slowed growth or a non-replicating persistent (NRP) state. NRP bacteria are tolerant to immune and antibiotic-mediated killing^7–9^, therefore understanding mechanisms underlying NRP may promote new methods to shorten the course of TB therapy.

Following macrophage infection, Mtb senses the mildly acidic pH of the phagosome and broadly remodels its gene expression^10^. Adaptations to acidic pH include the induction of the PhoPR regulon, induction of ESX-1 secretion, and remodeling of central metabolism and cell envelope lipids^11^. Defects in adaptations to acidic pH reduce Mtb virulence in macrophages and animals^12–15^, therefore, pH dependent adaptations are required for Mtb virulence.

Previous studies conducted by our lab sought to understand the interplay of acidic pH and Mtb central metabolism. We observed that Mtb exhibits selectivity of the carbon sources on which it can growth at pH 5.7 relative to pH 7.0. For example, Mtb incubated at acidic pH with glycerol as a sole carbon source is restricted for growth and establishes a viable, metabolically active state of NRP called acid growth arrest^16–18^. Acid growth arrest is observed on a variety of other carbon sources associated with glycolysis and TCA cycle. Interestingly, Mtb can resuscitate its growth at acidic pH by addition of specific carbon sources, such as pyruvate, acetate, oxaloacetate [OA] and cholesterol, which function at the intersection of glycolysis and the TCA cycle (a.k.a. the anaplerotic node)^18^. This discovery suggests that the anaplerotic node is the location of a pH-dependent metabolic switch that may promote Mtb growth on permissive carbon sources during pathogenesis at acidic pH, and that metabolic remodeling is required for pH adaptation^11^.

It is puzzling that Mtb cannot grow at acidic pH on specific carbon sources, as Mtb is provided with oxygen as a terminal electron acceptor and a carbon source that is well utilized at pH 7.0. Thus, acid growth arrest is different from other NRP models, where the bacterium is missing a key factor required for growth (*e.g.* oxygen or nutrients in the hypoxia or starvation models of NRP). Therefore, we hypothesized that acid growth arrest is not an actual restriction on growth, but an adaption by the bacterium to slow and arrest its growth. In a previous study, our lab previously sought to identify genes regulating acid growth arrest by selecting for mutants incapable of arresting their growth on minimal medium agar plates, buffered to pH 5.7 with glycerol as the carbon source. From our screening efforts, novel missense mutations were identified in *ppe51* (H37Rv annotated Mtb gene, *Rv3136*) and were named enhanced acid growth (*eag*) mutants ^16^. PPE51 is part of the PE and PPE mycobacterial protein family. Named for their unique N-terminus motifs Pro-Glu (PE) and Pro-Pro-Glu (PPE), most of these proteins have remained largely enigmatic in their functional roles^19,20^. However, a growing body of literature in recent years has assigned diverse putative functional roles for PE and PPE proteins including immune evasion^20–24^, calcium binding^25^, iron utilization^26,27^, Mg^2+^ and PO_3_^2−^ transport^28^, fibronectin binding^29^, and lipase activity^30,31^.

Mounting evidence suggests that some PE and PPE proteins may play important roles in Mtb nutrient acquisition. Examination of *pe* and *ppe* evolution reveals an expansion of this protein family corresponding with Type VII or ESX secretion systems, where it is thought that ancestral *pe* and *ppe* genes inserted into an *esx* gene cluster and expanded alongside this secretion system during subsequent gene duplication events ^22,32,33^. Secretion via ESX provides a route for PE and PPE proteins to access the cell surface and nutrients in the host cell milieu, which is supported by high-throughput proteomic evidence showing direct surface localization of PE and PPE proteins^26,28,34–36^. Mtb contains five ESX secretion systems^37^, with ESX-5 contributing to the majority of PE and PPE secretion^32,34,38^. Furthermore, ESX-5 and its cognate PE and PPE proteins have been implicated in the uptake of fatty acids and possibly the utilization of other nutrient substrates^34^. ESX-3-mediated PE and PPE proteins are thought to play a role in iron acquisition, whereby they have been shown to be directly involved in mycobactin-mediated iron uptake and heme uptake^26,39,40^. Taken together, these results provide direct examples of Mtb acquiring and utilizing host resources through secretion of PE and PPE proteins.

Based on these findings showing a role for PPE proteins in transport and that PPE51 *eag* variants could grow on glycerol, we hypothesized in our 2018 study^16^, “that these amino acid substitutions may increase Mtb growth by modulating mycomembrane permeability, possibly by modulating the channel size or specificity of PPE51, which may function as a porin to enhance access to glycerol or other nutrients at acidic pH.” The goal of this study was to test this hypothesis and further define the role of PPE51 in glycerol acquisition and pathogenesis. Notably, concurrent with this study, recently published studies confirmed the hypothesis that PPE51 is an exported cell surface-associated protein and linked to the nutrient acquisition of glycolytic carbon sources ^28,41,42^. Here, we show that in a saturating forward genetic screen only three PPE51 variants, S211R, E215K and A228D were isolated as *eag* mutants. The PPE51 variants specifically promoted growth at pH 5.7 on glycerol and not other tested carbon sources, supporting the substitutions selectively promote glycerol utilization. Radiolabeling studies show that the S211R variant has enhanced uptake of glycerol and accumulation of triacylglycerol (TAG), showing that the variants promote glycerol uptake. Differential thermal stability of WT versus S211R variant PPE51 proteins in the presences of glycerol, support the variant has direct and differential interactions with glycerol, but similar interactions with the non-permissive substrate glucose. Structural modeling supports that PPE51 forms a structure homologous with bacterial nutrient transporters, with the variants altering the predicted ligand specificity. These data are consistent with a model where PPE51 promotes uptake of glycerol across the mycomembrane by acting like a porin and that variants alter the conformation to enhance glycerol uptake. *eag* variants exhibit enhanced replication and reduced virulence in activated macrophages, supporting a role for pH-dependent slowed growth during macrophage pathogenesis.

## Methods

### Bacterial Strains and Growth Conditions

All experiments were performed with *M. tuberculosis* strains Erdman and CDC1551. Mtb was grown at 37 °C and 5% CO_2_ in vented T-25 culture flasks containing Middlebrook 7H9 media with 10% oleic acid-albumin-dextrose-catalase (OADC), 0.05% Tween-80, and 0.2% glycerol. For acid stress and single carbon source experiments, MMAT defined minimal media was used as described by Lee *et al*. ^43^: 1 g/L KH_2_PO_4_, 2.5 g/L Na_2_PO_4_, 0.5 g/L (NH_4_)_2_SO_4_, 0.17 g/L L-Asparagine monohydrate, 10 mg/L MgSO_4_, 50 mg/L ferric ammonium citrate, 0.1 mg/L ZnSO_4_, 0.5 mg/L CaCl_2_, and 0.05% Tyloxapol. MMAT media was buffered with 100 mM 3-(N-morpholino)propanesulfonic acid (MOPS) for experiments requiring pH 6.6-7.0 and 100 mM 2-(N-morpholino)ethanesulfonic acid (MES) for experiments requiring pH 5.5-6.5 ^44^. For growth curve experiments, Mtb was grown to mid-late log phase (OD_600_ 0.6-1.0) and seeded in MMAT at a starting OD_600_ of 0.05. Optical density measurements were conducted by removing 500 µL of samples at each time point. Viability assays were performed in a similar manner with samples being diluted 10-fold in PBS + 0.05% Tween-80 and plated for viable colony forming units (CFUs) on 7H10 + 10% OADC agar plates.

### Genetic Screen and Sequencing

A wild type Erdman Mtb population of 4×10^9^ bacteria was plated on MMAT agar plates (1 g/L KH_2_PO_4_, 2.5 g/L Na_2_PO_4_, 0.5 g/L (NH_4_)_2_SO_4_, 0.17 g/L L-Asparagine monohydrate, 10 mg/L MgSO_4_, 50 mg/L ferric ammonium citrate, 0.1 mg/L ZnSO_4_, 0.5 mg/L CaCl_2_, and 0.05% Tyloxapol) supplemented with 10 mM glycerol as the sole carbon source and buffered to pH 5.7 with 100 mM MES ^44^. Plates were incubated at 37 °C with spontaneous mutants appearing around week eight and isolated for growth. Single-colony isolates were confirmed as enhanced acid growth (*eag*) mutants under acidic conditions in liquid MMAT (pH 5.7) media amended with 10 mM glycerol. Whole genome sequencing (WGS) was performed on genomic DNA isolated from mutants representing various levels of enhanced growth as well as a wild type Erdman control. Samples were sequenced using the Illumina MiSeq in a 2×250-bp paired end format. Base calling was done by Illumina Real Time Analysis v1.18.54, demultiplexed, and converted to FastQ using Illumina Bcl2fastq v2.19.1. Low-quality bases were trimmed and adapter sequences were removed using Trimmomatic (v0.36)^45^ and aligned sequence reads to the Erdman reference genome using BWA-MEM^46^. SNPs and indels were identified using Genome Analysis ToolKit (GATK)^47^.

### Generation and Analysis of ppe51 Knockout

The *ppe51* gene was replaced with a hygromycin resistance (Hyg^R^) cassette in both Erdman and CDC1551 Mtb strain backgrounds using a new chromosomal engineering system called ORBIT (for “<u>o</u>ligonucleotide-mediated recombineering followed by <u>B</u>xb1 integrase targeting”) that combines site-specific recombination with homologous recombination^48^. An ORBIT recombineering plasmid (pKM444) expressing RecT annealase and Bxb1 integrase from the anhydrotetracycline (ATc)-inducible Ptet promoter and containing a kanamycin resistance (Kan^R^) cassette was first transformed into Mtb, selected for KanR, induced with ATc, and generated into electrocompetent cells. Electrocompetent cells were then transformed with a knockout integration plasmid (pKM464) harboring Hyg^R^ and targeting oligonucleotide. Hygromycin-resistant colonies were isolated and cured of the kanamycin-containing recombineering plasmid. Genomic DNA was extracted from transformants and the 5’ and 3’ junction sites of the knockout were confirmed by PCR and sequencing using ORBIT target-specific and *ppe51*-specific primers (Figure S7B-C, Table SX). Gene replacement was further verified via quantitative real time PCR (qPCR) (Figure S7D). *Δppe51* was complemented with WT and variant *ppe51* from their native promoter and confirmed by qPCR (Figure S7E).

### pH-and-Glycerol dose response combination growth assays

Mtb cultures were incubated in a range of pH buffered MMAT media (pH 5.0-pH 7.0) at a starting OD_600_ of 0.2 in 96-well plates. Cultures were treated with 2.5-fold dose-response (0.13-80 mM) of glycerol and incubated over the course of 21 days, with growth assessed by optical density. Bacterial viability was assessed by diluting wells 10-fold and plating for viable CFUs on 7H10 + 10% OADC agar plates. Optical density data was converted to percent of maximum well-growth and normalized based on no carbon control at pH 5.5 (0%) and maximum Mtb growth on glycerol at pH 6.5 (100%). Each condition and time-point experiment was conducted in triplicate and representative of multiple individual experiments.

### Radiolabeled Glycerol Uptake Assay

Mtb Erdman cultures were pre-adapted for 3 days in MMAT media (pH 5.7 or pH 7.0) containing 10 mM glycerol. Following adaptation, Mtb was washed twice with PBS+0.05% Tween-80 and resuspended in the same buffered MMAT media amended with 10 mM glycerol and 6 μCi of [U-^14^C] Glycerol. Samples were removed over the course of 24 hours, fixed with 4% paraformaldehyde, and assessed for total radioactivity using scintillation counting. Short-term radiolabeling uptake experiments were also conducted in a similar manner as previously described^28,49^ to compare previously published results. Mtb strains were pre-adapted for 3 days in MMAT media buffered to pH 5.7 with 10 mM glycerol, harvested via centrifugation, and washed twice with uptake buffer (50 mM Tris, pH 7.0, 15 mM KCl, 10 mM (NH4)2SO4 1mM MgSO4, 0.02% tyloxapol). Samples were resuspended in uptake buffer at a starting OD_600_ of 2.0 and incubated for 30 minutes at 37 °C in T-25 tissue culture flasks. Cultures were amended with 16 µM [U-^14^C] Glycerol, and 200 µL samples were taken at minutes 3, 6, 9, 18, and 36. Samples were immediately filtered by centrifugation through a 0.22 µM Spin-X tube (Costar) and washed twice with 4% paraformaldehyde. Total radioactivity of the filters was assessed by liquid scintillation counting. All strains used for radiolabel uptake experiments were repeated in two biological replicates.

### Analysis of metabolism of radiolabeled lipids into Mtb lipids

Mtb Erdman cultures were pre-adapted as described above for the uptake experiments. Following pre-adaptation, cultures were seeded at a starting OD_600_ of 0.2 in MMAT media (pH 5.7 or pH 7.0) + 10 mM glycerol and set up in two biological replicates. Lipids were labeled with 6 μCi of [U-^14^C] Glycerol for 6 days, and samples were pelleted and washed with PBS before lipid extraction. Total lipids were extracted and Folch washed as previously described^18^ and ^14^C-incorporation was measured using scintillation counting. For thin layer chromatography (TLC), 5,000 counts per minute (CPM) of each pH 5.7 sample and 10,000 CPM of each pH 7.0 sample was loaded on a 100-cm^2^ high-performance TLC silica gel 60 aluminum sheet (EMD Millipore) and analyzed with a chloroform:methanol:water (90:10:1 v/v/v) solvent system^50^. Sulfolipids, TAG, and PDIM were separated as previously described^13,18,50^ and quantified using a phosphor screen and Typhon imager and ImageJ software^51^.

### Replication during acid growth arrest

For measurement of replication during acid growth arrest, WT Mtb, *Δppe51*, and *ppe51* native variants in both CDC1551 and Erdman backgrounds carrying the pBP10 plasmid^52^ were inoculated into MMAT media (pH 5.7 and pH 7.0) + 10 mM glycerol and in the absence of kanamycin. Plasmid loss and percentage of bacteria still containing the pBP10 was determined by plating for CFUs on 7H10 + 10% OADC agar plates ±25 µg/uL kanamycin. Rates of growth, death, and cumulative bacterial burden were quantified using equations as previously described^53^. The Mtb segregation constant (s = 0.18± 0.023) was used for all calculations.

### Macrophage pathogenesis studies

Bone Marrow-derived macrophages (BMDMs) were extracted and infected with the panel of complemented strains built into the CDC1551 *Δppe51* knockout background at a multiplicity of infection (MOI) of 1:1 using previously described methods^54^. BMDMs were activated by treating with 100 units/mL IFN-γ overnight, followed by treatment with 10 ng/mL lipopolysaccharide overnight. Infected BMDMs were lysed at days 0, 3, 6, and 9 and intracellular bacterial lysates were serially diluted and enumerated on 7H10 + 10% OADC agar plates. Each strain for each timepoint was performed in triplicate. BMDMs were also infected with CDC1551 strains containing the pBP10 replication clock plasmid as described in the pBP10 *in vitro* experiments using the same macrophage infection methods described above with minor modifications. BMDMs infected with pBP10-containing strains were lysed at days 0, 2, 4, 6, and 8 and enumerated on 7H10 + 10% OADC agar plates ±25 µg/uL kanamycin selection. Calculations for pBP10 plasmid loss and replication dynamics were performed as described in the *in vitro* pBP10 experiments.

### Recombinant PPE51 Protein Expression and Purification

The ORF of PPE51 was amplified using pET23::*ppe51*_FWD and pET23::*ppe51*_REV primers (Table S1) and cloned into the pET23a+ vector containing a C-terminal polyhistidine (His)-tag. The cloned protein has a deletion of the final four C-terminal amino acids. Transformants propagated in *E. coli* BL21(DE3) were selected on LB agar plates containing Amp. The S211R mutation was introduced into the pET::*ppe51*-WT construct using the site-directed mutagenesis primers PPE51-S211R_FWD and PPE51-S211R_REV (Table S1) and the QuikChange^TM^ Site-Directed Mutagenesis Kit. Overnight cultures were expanded into 250 mL of fresh LB media with Amp at an initial inoculum OD_600_ of 0.05 and grown to an OD_600_ of 0.6 at 37 °C with shaking at 200 rpm. Proteins were then induced with 1 mM of isopropyl-ß-D-thiogalactopyranoside (IPTG) at 18°C for 20 hours. Culture was then harvested via centrifugation at 4000 rpm for 25 minutes at 4°C. Pellets were then lysed for 30 minutes on ice with occasional vortexing in ice-cold lysis buffer (50 mM phosphate buffer [pH 7.6], 200 mM NaCl, 0.1% Triton-x-100, 0.1 mg/mL PMSF, 0.5 mg/mL lysozyme). Because PPE51 possibly interacts with glycerol, glycerol was completely removed from all buffers used during the purification process. Cells were further lysed by sonication and cell lysate was clarified via centrifugation at 14,000 rpm for 30 minutes at 4°C. Supernatant was loaded onto a nickel ion-containing affinity resin column and bound overnight with shaking at 4°C. Protein was washed first with wash buffer containing no imidazole and a second time with wash buffer containing 50 mM imidazole. PPE51 protein was then eluted into (4) 1 mL fractions with elution buffer containing 200 mM imidazole. Recombinant PPE51 was quantified using the Qubit assay.

### PPE51 Protein Thermostability Assay

The thermostability assay was performed as previously described^55^ with 13.5 µL of 0.635 mg/mL of batch-purified PPE51 samples aliquoted into PCR tube containing 1.5 µL of 100 mM glycerol, yielding a final glycerol concentration of 10 mM. Samples were incubated for 20 minutes at room temperature and transferred to PCR thermocyclers where they were incubated for an additional 5 minutes at the following temperatures: 35, 40, 45, 50, 55, 60, and 65°C. Samples were then centrifuged at 4000 rpm for 10 minutes to pellet precipitated protein. After centrifugation, soluble protein was removed from the tubes and detected in Western blots using mouse anti-His tag monoclonal antibody followed by HRP-conjugated anti-mouse IgG secondary antibody. Enhanced chemiluminescence (ECL) Western blotting substrate (Pierce) was used for Western blot detection. The AI600 Chemiluminescent Imager was used to visualize and analyze Western blot results.

## Results

### All isolated eag mutants have spontaneous mutations in ppe51

During acid growth arrest in minimal media, Mtb is provided all required nutrients including a metabolically utilized carbon source and a terminal electron acceptor. This suggests that acid growth arrest is not due to a physiological limitation presented by the growth environment but rather is a regulated process whereby Mtb adapts to its acidic environment. A previously published forward genetic screen tested this hypothesis using a CDC1551 transposon mutant library containing >100,000 was plated on MMAT defined minimal media agar with glycerol at pH 5.7 and resulted in 98 transposon (Tn) mutants and two spontaneous WT mutants^16^. These mutants were isolated and confirmed as *<u>e</u>nhanced <u>a</u>cid <u>g</u>rowth* (*eag*) mutants based on their ability to grow well compared to native WT Mtb at pH 5.7 in liquid MMAT supplemented with glycerol ^16^. Interestingly, complementation attempts with the Tn mutants did not restore growth arrest, and whole genome sequencing identified spontaneous mutations in *ppe51* in both Tn and WT mutant backgrounds ^16^. To repeat a saturating screen, in the absence of transposon mutagenesis, a second forward genetic screen was performed on MMAT agar buffered to pH 5.7 with glycerol, using a larger bacterial population (4×10^9^ bacteria) in the Erdman Mtb strain (Figure 1A). From the WT Erdman screen, 98 spontaneous *eag* mutants were isolated of which 52 were colony-purified and confirmed for enhanced growth under acidic conditions in liquid MMAT containing glycerol (Figure 1B, Figure S1A-D). The *eag* isolates exhibited an up to ∼4-fold increase in growth compared to WT Erdman which exhibited complete growth arrest (Figure 1B). Of these *eag* mutants, 24 were selected for whole genome sequencing. Remarkably, all 24 isolates had single nucleotide polymorphisms (SNPs) mapping to the *ppe51* gene (Table 1). All mutations were non-synonymous (S211R, A228D, and E215K) and were centrally located within a 50 bp region on the *ppe51* gene (Figure S2). The S211R and A228D variants were also identified in the prior Tn mutant CDC1551 screen, with E215K being a novel mutation found in the new Erdman screen.

**Figure 1.**
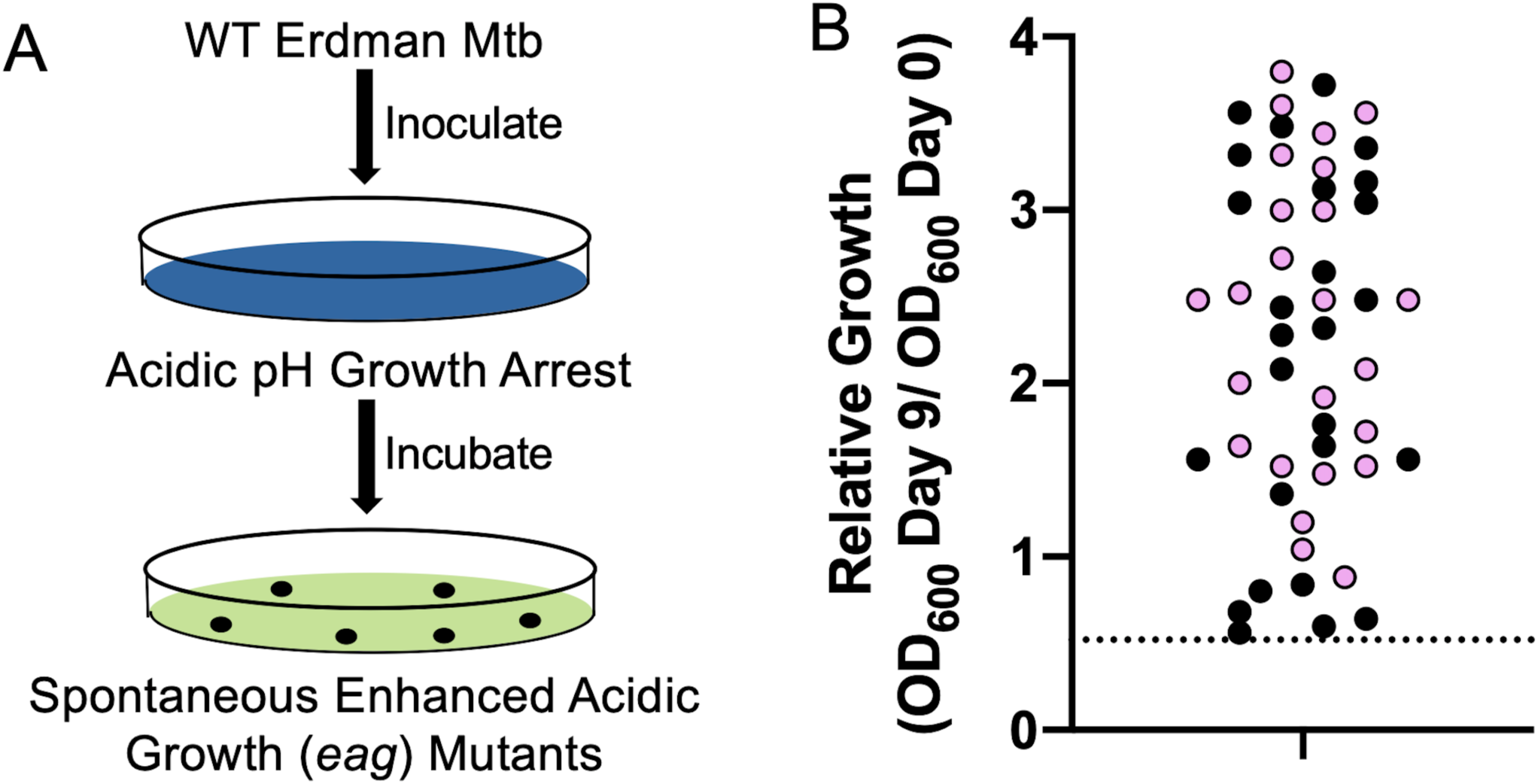
Genetic screen for mutants unable to arrest growth at acidic pH. **A)** Schematic of the forward genetic screen used to acquire *eag* mutants. The appearance of distinct colony growth after 8 weeks was indicative of mutants that were unable to arrest their growth at acidic pH. **B)** Growth phenotypes of isolated mutants were determined by measuring Day 9 OD_600_ and compared to OD_600_ from the initial inoculum. Each dot represents mutants that were isolated from MMAT agar plates and confirmed for enhanced growth in liquid MMAT (pH 5.7). Pink-colored data points indicate mutants chosen for whole-genome sequencing. The dotted line represents relative WT Erdman growth (< ratio of 1).

**Table 1.** Whole genome sequencing results of isolated eag variants

| Plate No. | Mutant # | SNP Location | Nucleotide Change | Amino Acid Change | Quality Score |
| --- | --- | --- | --- | --- | --- |
| Plate 1 | *eag1.7* | 3497961 | G<u>C</u>C→G<u>A</u>C | A228D | 5478 |
|  | *eag1.8* | 3497911 | AG<u>C</u>→AG<u>A</u> | S211R | 6929 |
|  | *eag1.12* | 3497961 | G<u>C</u>C→G<u>A</u>C | A228D | 7373 |
|  | *eag1.14* | 3497911 | AG<u>C</u>→AG<u>A</u> | S211R | 6328 |
|  | *eag1.33* | 3497911 | AG<u>C</u>→AG<u>A</u> | S211R | 5270 |
|  | *eag1.34* | 3497911 | AG<u>C</u>→AG<u>A</u> | S211R | 5270 |
| Plate 2 | *eag2.1* | 3497961 | G<u>C</u>C→G<u>A</u>C | A228D | 4417 |
|  | *eag2.2* | 3497961 | G<u>C</u>C→G<u>A</u>C | A228D | 6803 |
|  | *eag2.3* | 3497911 | AG<u>C</u>→AG<u>A</u> | S211R | 5205 |
|  | *eag2.6* | 3497911 | AG<u>C</u>→AG<u>A</u> | S211R | 5976 |
|  | *eag2.8* | 3497961 | G<u>C</u>C→G<u>A</u>C | A228D | 4270 |
|  | *eag2.14* | 3497921 | <u>G</u>AG→<u>A</u>AG | E215K | 3442 |
|  | *eag2.16* | 3497911 | AG<u>C</u>→AG<u>G</u> | S211R | 4575 |
| Plate 3 | *eag3.2* | 3497961 | G<u>C</u>C→G<u>A</u>C | A228D | 4140 |
|  | *eag3.4* | 3497961 | G<u>C</u>C→G<u>A</u>C | A228D | 3292 |
|  | *eag3.9* | 3497911 | AG<u>C</u>→AG<u>G</u> | S211R | 3645 |
|  | *eag3.15* | 3497911 | AG<u>C</u>→AG<u>A</u> | S211R | 3784 |
|  | *eag3.23* | 3497911 | AG<u>C</u>→AG<u>G</u> | S211R | 4609 |
| Plate 4 | *eag4.5* | 3497961 | G<u>C</u>C→G<u>A</u>C | A228D | 5153 |
|  | *eag4.7* | 3497911 | AG<u>C</u>→AG<u>G</u> | S211R | 5107 |
|  | *eag4.12* | 3497911 | AG<u>C</u>→AG<u>G</u> | S211R | 5179 |
|  | *eag4.21* | 3497961 | G<u>C</u>C→G<u>A</u>C | A228D | 3958 |
|  | *eag4.24* | 3497911 | AG<u>C</u>→AG<u>G</u> | S211R | 5517 |

### ppe51 mutations are sufficient to overcome growth arrest

Given that *ppe51* variants exhibit enhanced growth at acidic pH, we investigated the function of the variant alleles in the presence of the WT *ppe51* allele. Overexpression constructs of WT or mutant *ppe51* were transformed into WT CDC1551 or WT Erdman Mtb strains carrying the native *ppe51* allele. Overexpression strains were grown in MMAT at pH 5.7 with glycerol as a carbon source. Overexpression of *ppe5*1-S211R and *ppe51*-A228D variants in WT Mtb resulted in significantly enhanced growth under acidic conditions (Figure 2A). In contrast, overexpression of WT *ppe51* and the empty vector exhibited complete growth arrest at pH 5.7. The E215K allele was also not sufficient at overcoming growth arrest which may explain why it has only been observed once across two independent forward genetic screens. Additionally, all overexpression strains grew equally well at pH 7.0 (Figure S3), showing that the observed growth phenotype is pH-specific. Interestingly, although the growth phenotype with A228D overexpression resulted in enhanced growth at acidic pH, it grew at a slower rate compared to S211R overexpression strains in both CDC1551 and Erdman backgrounds. We examined individual variant alleles from the screen (Figure 1B) and observed that S211R variants significantly grouped together at a higher rate of growth compared to A228D and E215K (Figure 2B and 2C). Together, these results demonstrate that the *eag* mutations confer a dominant, gain-of-function growth phenotype, and specific mutations are associated with differential strength of the phenotype.

**Figure 2.**
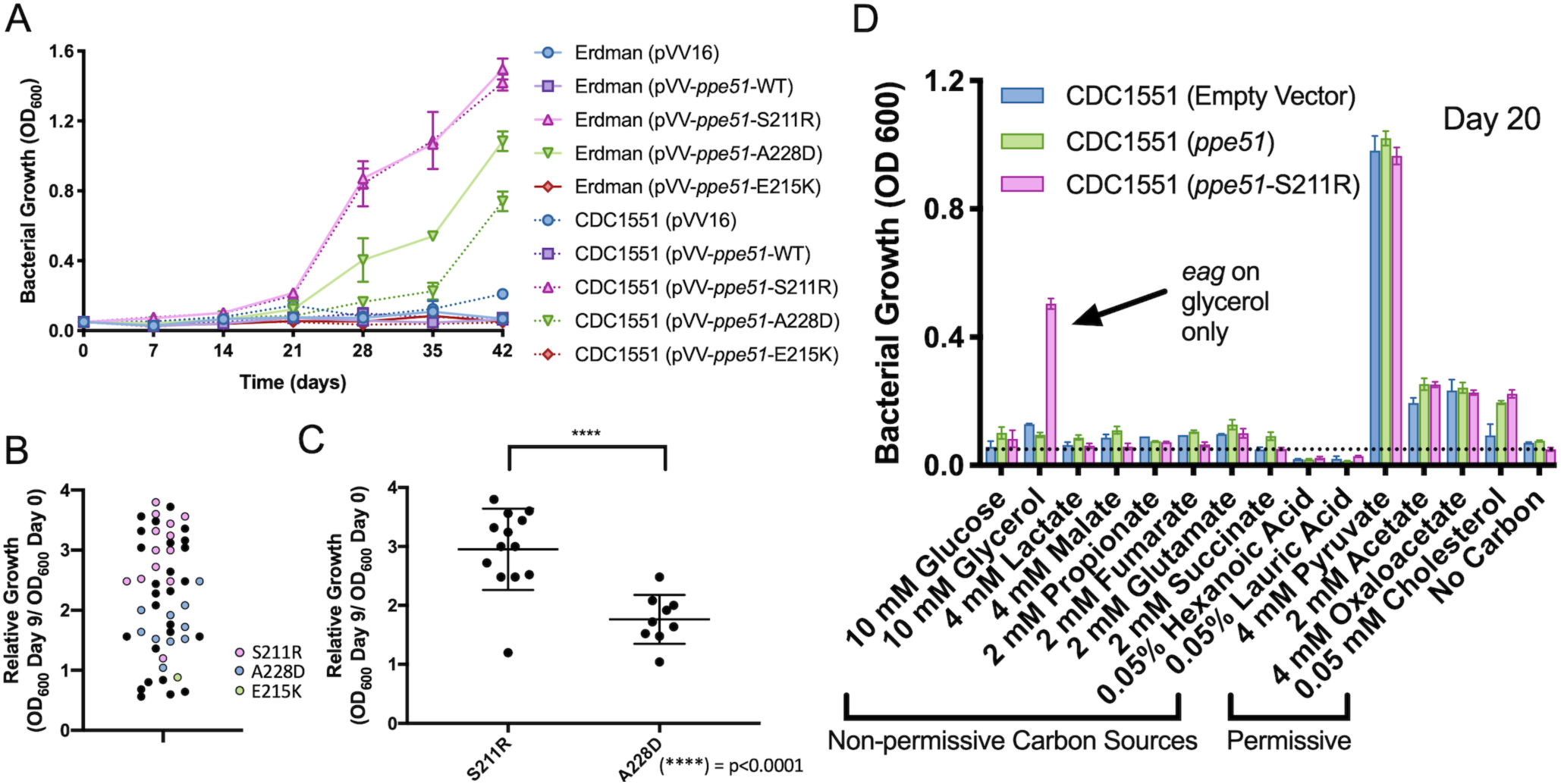
PPE51 variants drive the *eag* phenotype and exhibit phenotypic and carbon source-dependent growth differences. **A)** Growth curve of WT Mtb (Erdman and CDC1551 strains) overexpressing *ppe51* and *eag* variants. Growth of WT overexpression strains (pVV16 [empty vector] and pVV-*ppe51*-WT) were compared to WT carrying the overexpression constructs of the mutant alleles (S211R, A228D, and E215K) in minimal media (pH 5.7 + glycerol). Overexpression of *eag* mutant alleles in WT Mtb results in significantly enhanced growth under acidic conditions. **B)** *eag* variants show distinct clustering of variant type based on relative growth. **C)** Statistical analysis of growth differences between native *eag* strains containing S211R or A228D was performed using an unpaired t-test (****P < 0.0001) with Welch’s correction. **D)** Analysis of the S211R variant growth on various carbon sources. CDC1551 overexpression strains were grown in MMAT medium (pH 5.7) in the presence of various growth-permissive (i.e. pyruvate acetate, OA) and non-permissive (i.e. glucose, propionate, lactate) carbon sources. *ppe51*-S211R (pink bars) growth is carbon source specific and only exhibits enhanced growth on glycerol, a normally non-permissive carbon source at pH 5.7. Growth on permissive carbon sources is not impacted by *ppe51*-S211R. The horizontal dotted line indicates the starting density of 0.05 OD_600_. Similar results were observed in Erdman (Figure S5).

### PPE51 variants selectively promote growth on glycerol

Based on the enhanced growth phenotype that the variants exhibit at acidic pH, we hypothesized that this phenotype may be due to *ppe51* variants modulating mycomembrane permeability, resulting in enhanced nutrient uptake. To test this hypothesis, we conducted an Ethidium Bromide (EtBr) assay looking at permeability with WT Mtb overexpression constructs (empty vector, WT, and S211R) in both CDC1551 and Erdman backgrounds. With the EtBr assay we did not observe differences in the rate of uptake between the WT and *eag* overexpression strains in either CDC1551 or Erdman (Figure S4), suggesting that the growth phenotype is not due to a general increase in permeability. We then hypothesized that the growth phenotype may be due to nutrient-specific uptake. We explored this possibility by growing CDC1551 and Erdman overexpression strains (empty vector, *ppe51* and *ppe51*-S211R) in liquid minimal media (pH 5.7) in the presence of various growth-permissive (*e.g.* pyruvate, acetate, cholesterol) and non-permissive (*e.g.* glucose, glycerol, propionate) carbon sources^18^. After 20 days we found that enhanced acid growth was only observed with *ppe51*-S211R in the presence of 10 mM glycerol, a normally non-permissive carbon source at pH 5.7, in both CDC1551 and Erdman (Figure 2D, Figure S5, Figure S6A-B). As expected, all overexpression strains exhibited enhanced growth on permissive carbon sources in both Mtb backgrounds. Notably, the *ppe51* variants specifically promote growth on glycerol, but not glucose, another proposed nutrient associated with PPE51-dependent uptake^28,41^, demonstrating the variants are selective for glycerol.

### ppe51 is not required for survival during acid growth arrest

Transcriptional profiling studies previously conducted show that *ppe51* is significantly induced during acid growth arrest^18^.We hypothesized that *ppe51* may be required for Mtb to promote survival when exposed to acid growth arresting conditions. To test this hypothesis, a Δ*ppe51* knockout strain was generated in both CDC1551 and Erdman Mtb using the mycobacteria-specific ORBIT system (Figure S7A)^48^. Successful knockout of *ppe51* was confirmed by sequencing the *oriE* and HygC junction sites of Δ*ppe51*, PCR amplification of the entire knockout region, and RT-PCR (Figure S7B-D). Complementation constructs containing the native *ppe51* promoter were introduced into Δ*ppe51* carrying WT *ppe51*, variant *ppe51* (S211R, A228D, E215K) and a double variant (S211R+A228D). An empty complementation vector was also introduced into Δ*ppe51.* The presence of these constructs was also confirmed via RT-PCR (Figure S7E). Growth curves of the complementation constructs grown in growth arresting conditions showed that WT and empty vector strains exhibit growth arrest at pH 5.7, whereas the variant complemented strains exhibited enhanced growth in both CDC and Erdman (Figure 3A, Figure S8A). Additionally, the S211R, A228D, and S211R+A228D variant complemented strains grow slightly better compared to E215K in both CDC1551 and Erdman Mtb strain knockout backgrounds, which aligns with previous overexpression growth curve and relative growth data (Figure 2A and B). At pH 7.0, all WT and complemented strains showed similar levels of growth (Figure S8B and C). To examine the role of *ppe51* in survival, a viability assay was performed with the previously described panel of strains. The empty vector and WT-complemented Δ*ppe51* maintain viability over the course of 40 days at pH 5.7 (Figure 3B, Figure S9A). The complemented *ppe51* variants also maintain viability and replicate at pH 5.7. At pH 7.0, all WT and complemented strains maintain viability and exhibit similar increases in CFUs over the course of 18 days (Figure S9B and C).

**Figure 3.**
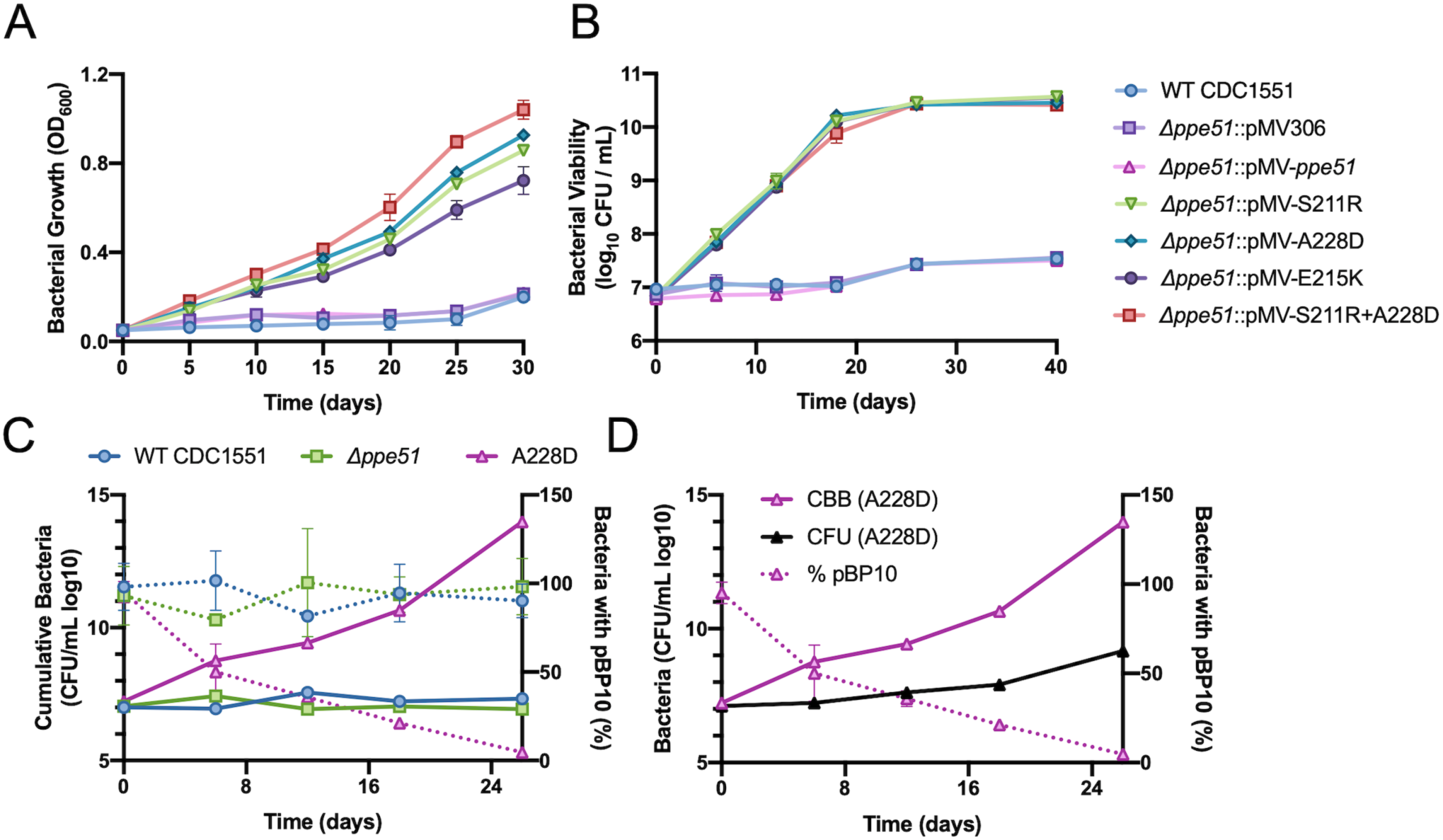
Viability and replication dynamics of *eag* mutants. **A)** Growth curve of *Δppe51* complemented from its native promoter with the WT *ppe51* allele or mutant alleles and performed under acid growth arrest conditions: minimal media buffered to pH 5.7 with glycerol as a carbon source. *eag* variants promote Mtb growth at pH 5.7, while *Δppe51* complemented with WT *ppe51* maintains growth arrest. **B)** Viability of Mtb strains as measured by colony-forming units (CFUs). **C)** CDC1551 Mtb containing the native variant allele, A228D, continues to replicate during acid growth arrest conditions, but WT Mtb and *Δppe51* cease replication. To estimate replication dynamics of the indicated strains, clock plasmid replication data was obtained from CFU counts (right axis, dotted lines). CFUs of plasmid-free and plasmid-bearing strains were then used to calculate cumulative bacterial burden (CBB) of total live, dead, or degraded Mtb (left axis, solid lines). **D)** Replication dynamics of the native A228D (CDC1551) variant, comparing CBB (cumulative bacterial burden), CFU (total CFUs from nonselective plating), and % pBP10 (percentage of bacteria carrying plasmid).

The stable viability of WT or Δ*ppe51* mutant may be due to growth arrest or, alternatively, balanced growth and death. To determine if the strains are truly growth arrested we examined replication using a the pBP10 clock plasmid, transformed into WT Mtb, Δ*ppe51*, and *eag* variants in both CDC1551 and Erdman backgrounds^53^. The strains were incubated in minimal media (pH 5.7 and 7.0) with glycerol for 40 adays. We observed that the WT and Δ*ppe51* strains do not replicate under acid growth arrest conditions in both strain backgrounds (Figure 3C, Figure S10B). In contrast, we are able to observe high rates of replication in the *eag* variants at pH 5.7 (Figure 3C, Figure S10B). We then compared *eag* variants’ calculated cumulative bacterial burden (CBB) to total CFUs counted on nonselective plates and observed that greater rates of replication in the *eag* variants is associated with a high death rate, yielding a large difference between CBB and total CFUs (Figure 3D, Figure S10D-F). When *in vitro* pBP10 growth studies were conducted at pH 7.0 with all strains, we observed similar high rates of replication and plasmid loss across all strains (Figure S10A and C). Interestingly, these results show that enhanced growth at acidic pH is driven by higher replication, but this growth is offset somewhat by a higher death rate, supporting the conclusion that faster replication at acidic pH is deleterious to Mtb survival.

### Acidic pH limits glycerol uptake and PPE51 variants overcome this restriction

Pyruvate can rescue growth on glycerol in a concentration-dependent manner at pH 5.7 ^18^. However, it is unknown whether glycerol concentration affects acid growth arrest. We hypothesized that acid growth arrest may be driven by glycerol starvation and the PPE51 variants promote growth by promoting enhanced uptake of glycerol. If this is the case, we would expect to see a dependence of glycerol concentration and acidic pH on growth. To examine this, we examined checker-board dose responses combining varying pH levels (pH 6.5-5.5) and glycerol concentrations (80 mM-0.13 mM) using the panel of isogenic strains. The standard concentration of glycerol used in our acid growth arrest model is 10 mM. Growth in the wells was analyzed using optical density (OD_600_) and data was normalized to wells containing the highest (100%) levels of growth and wells with no carbon representing the lowest (0%) levels of growth. Growth assays were performed for 21 days, and the data shown is Day 14 which is representative for the duration of the experiment. Interestingly, we found that growth arrest appears to be both pH and glycerol concentration-dependent, with growth partially rescued at high concentrations of glycerol (∼80 mM) for WT, Δ*ppe51*::pMV306 and Δ*ppe51*::pMV-*ppe51* at pH 5.7 (Figure 4 and S11A). Additionally, we observed higher levels of growth at lower glycerol concentrations (∼0.82 mM) at pH 5.7 with the complemented *ppe51* variants compared to the empty vector and WT *ppe51* complemented strains. Growth could also be rescued with high glycerol concentration (∼32 mM) for variants at pH 5.5. Interestingly, the presence of the double *ppe51* variant (S211R+A228D) overcomes growth arrest at pH 5.5 at even lower glycerol concentrations (∼5.12 mM) compared to the single variants, indicating that the presence of two *eag* point mutations confers a slight growth advantage during acid growth arrest. Concentrations of glycerol below 0.33 mM do not rescue growth starting at pH 6.0 in any *eag* strains, which could be due to glycerol being fully consumed. Similarly, these observations were also made in the native *eag* variants in both CDC1551 and Erdman, while WT exhibited a reduced capacity for glycerol uptake (Figure S11B). Together, these findings suggest that Mtb has reduced capacity to uptake glycerol in a pH-dependent manner, and that PPE51 variants function by promoting enhanced uptake of glycerol.

**Figure 4.**
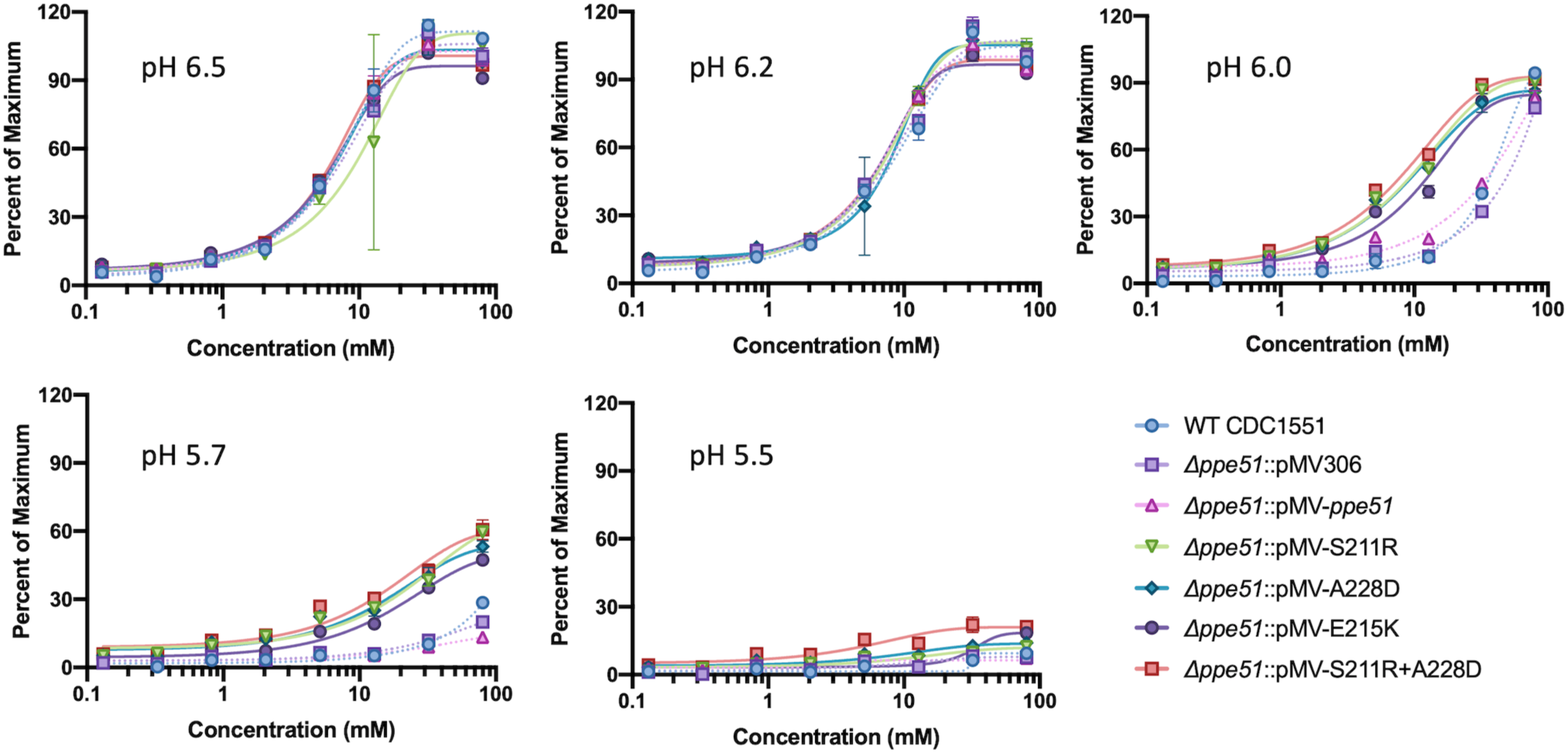
Mtb restricts glycerol uptake at low pH. Growth of WT CDC1551, *Δppe51* (empty vector), and *Δppe51* complemented strains in minimal media supplemented in a dose-dependent manner with glycerol and buffered to one of five pH levels (pH 6.5, 6.2, 6.0, 5.7, or 5.5). All strains exhibit a reduced capacity for growth starting ∼2 mM glycerol compared to higher glycerol concentrations. At decreasing pH, WT, *Δppe51*(empty vector), and *Δppe51*::pMV-WT restrict their ability to uptake glycerol, whereas any variant complement is able to maintain glycerol uptake. However, restricted growth can be rescued at high concentrations of glycerol (∼80 mM) at pH 5.7 for WT, *Δppe51*(empty vector), and *Δppe51*::pMV-*ppe51*, and pH 5.5 for variant complements. Growth analyses were performed at Day 14 following initial inoculation with data being shown as percent of the maximum well-growth. All conditions were conducted in triplicate and representative of multiple independent experiments. Similar data were observed in a native *eag* mutant (Figure S11B).

Based on these checkerboard results, *ppe51* appears to restrict its growth on glycerol at acidic pH. Additionally, WT Mtb has been shown to completely arrest its growth at pH 5.7 on 10 mM glycerol; however, it is able to maintain viability for up to 40 days, remains metabolically active, and incorporate limited amounts of exogenous ^14^C-glycerol into lipids^16^. To further test the hypothesis that Mtb restricts glycerol uptake at acidic pH and that *eag* variants promote enhanced glycerol uptake, a radiolabeling experiment using ^14^C-glycerol was conducted with WT Erdman and the Δ*ppe51* complemented strains previously described. Strains were pre-adapted for three days in MMAT (pH 5.7 or 7.0) with 10 mM glycerol and washed with PBS prior to radiolabeling with 6 µCi of ^14^C-glycerol. Samples were collected over the course of 24 hours, washed, and analyzed for radiolabel uptake by scintillation counting. All complemented strains containing a *ppe51* variant accumulated ^14^C-glycerol at a similarly increased rate of approximately 300% compared to the WT Mtb strain (Figure 5A). These results are consistent with radiolabeling that was conducted with the pVV16 overexpression empty vector, S211R overexpression strain, and native *eag*-S211R variant where we observed similar enhanced glycerol uptake at ∼ 60% with strains containing S211R compared to WT overexpression empty vector (Figure S12A). We also looked at glycerol uptake with WT CDC1551, empty vector, and complemented S211R at pH 7.0. We did not observe significant differences in glycerol uptake between strains, and the rate of uptake was similar to the complemented *ppe51* variant strains at pH 5.7 (Figure S12B). Together, these results show that Mtb does restrict glycerol uptake at pH 5.7 regardless of whether *ppe51* is functionally intact. In contrast, strains containing *ppe51* variants have significantly enhanced glycerol uptake.

**Figure 5.**
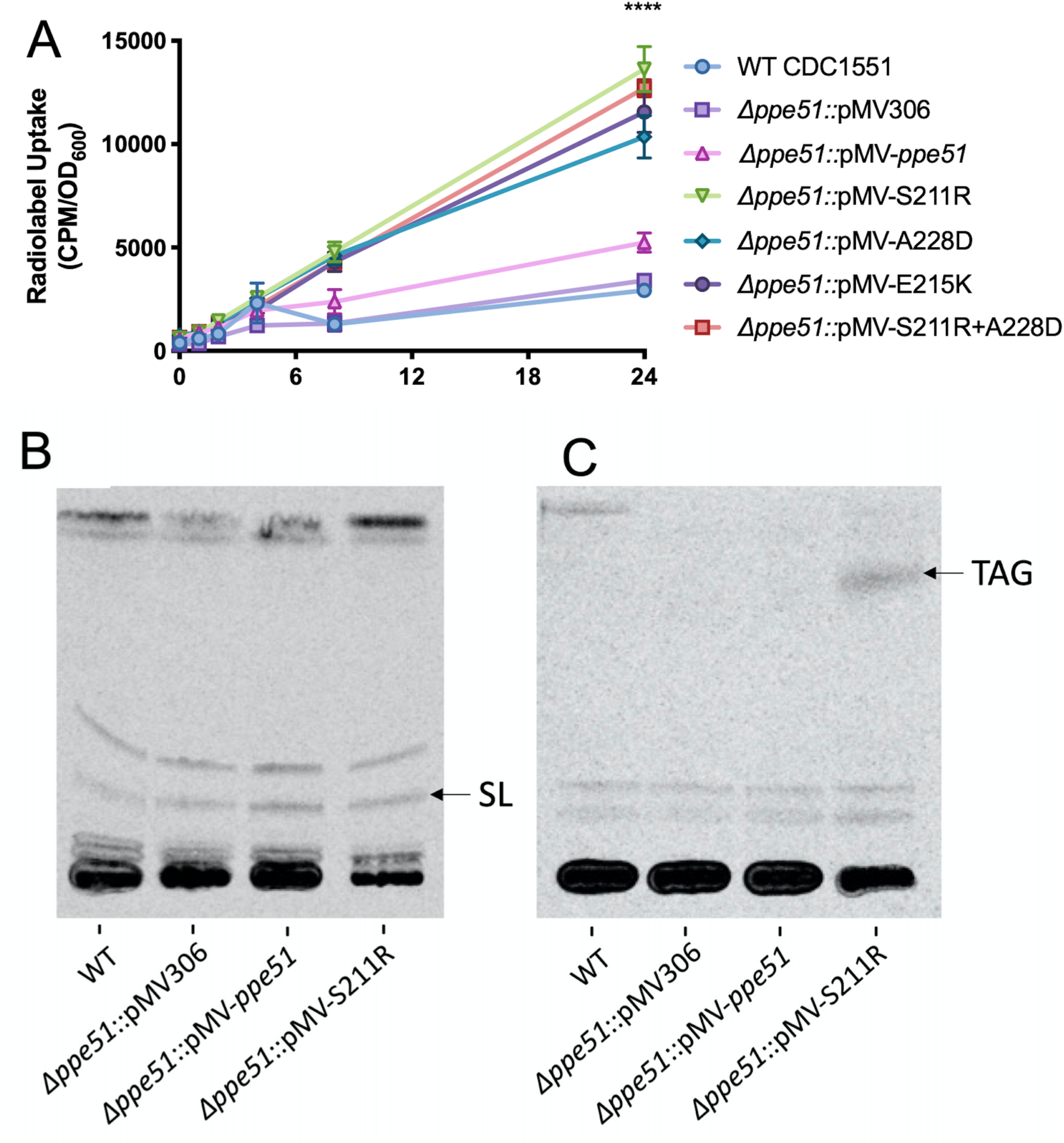
*eag* variants exhibit enhanced ^14^C-glycerol uptake and incorporation into lipids. **A)** Strains expressing *eag* variants uptake ^14^C-glycerol at an enhanced rate. Mtb was pre-adapted for 3 days in MMAT (pH 5.7) with 10 mM glycerol and subsequently washed prior to the addition of radiolabeled glycerol. ^14^C-glycerol uptake was measured using scintillation counting at various time-points over the course of 24 hours. Significance was determined by two-way ANOVA (Tukey’s multiple comparisons test; ****P < 0.0001) **B)** Incorporation of ^14^C-glycerol into sulfolipids at acidic pH. Sulfolipid is indicated with an arrow and accumulates at a similar rate in each strain. Strains were analyzed in duplicate with representative results being shown. **C)** Incorporation of ^14^C-glycerol into TAG at acidic pH. TAG are indicated with an arrow and are absent from all strains except for *Δppe51*::pMV-S211R. Strains were analyzed in duplicate with representative results being shown.

While the radiolabeling strongly indicated that glycerol was being taken up by the strains, it did not answer whether glycerol was being metabolized by Mtb and incorporated into lipids or binding to the mycomembrane without uptake across the plasma membrane. To address this question, we performed lipid radiolabeling with ^14^C-glycerol. WT Erdman and Δ*ppe51* complemented strains were pre-adapted for three days in the same culture conditions as the previously described radiolabeled uptake experiment. The operon controlling sulfolipid synthesis is induced in a *phoPR*-dependent and a pH-dependent manner^18,50,56^. Sulfolipid was observed to specifically accumulate at pH 5.7 (Figure 5B) with no accumulation occurring at pH 7.0 (Figure S12C). Triacylglycerol (TAG) has been shown to accumulate during periods of hypoxic and pH-stress^18,57^, and pathways involved in TAG synthesis play a role in reducing Mtb growth by redirecting carbon flux away from the TCA cycle^58^. Interestingly, we found that TAG accumulated specifically in the complemented S211R strain at pH 5.7 (Figure 5C). In contrast, we did see similar TAG accumulation across all strains at pH 7.0 (Figure S12D). The observation of labeled lipids in both growth arrested and growing Mtb at acidic pH, shows that glycerol is imported and metabolized at acidic pH, with enhanced uptake in the S211R variant.

### PPE51 variants have selectively reduced growth in activated macrophages

*ppe51* is induced in a pH-dependent and *phoP*-dependent manner within 2 hours following phagocytosis by macrophages^10^, suggesting that *ppe51* is important for pathogenesis. We hypothesized that *ppe51* or its *eag* variants may be required for pathogenesis, specifically in activated macrophages, where the phagosome is acidified. To test this hypothesis, resting and activated primary murine bone marrow-derived macrophages (BMDMs) were infected with WT CDC1551 and Δ*ppe51* mutant and complemented variant strains. In resting macrophages, we did not observe significant differences in Mtb growth between the strains (Figure 6A), with all of the strains growing ∼1.25-log over nine days. In contrast, in activated macrophages, while the WT, Δ*ppe51* empty vector, and Δ*ppe51* WT complemented strain still exhibit ∼1.25-log increase in growth, the *ppe51* complemented variants show significantly lower growth (Figure 6B). These results show that *ppe51* variants are selectively attenuated for virulence in activated macrophage environment, which is consistent with a pH-dependent phenotype that is observed *in vitro*.

**Figure 6.**
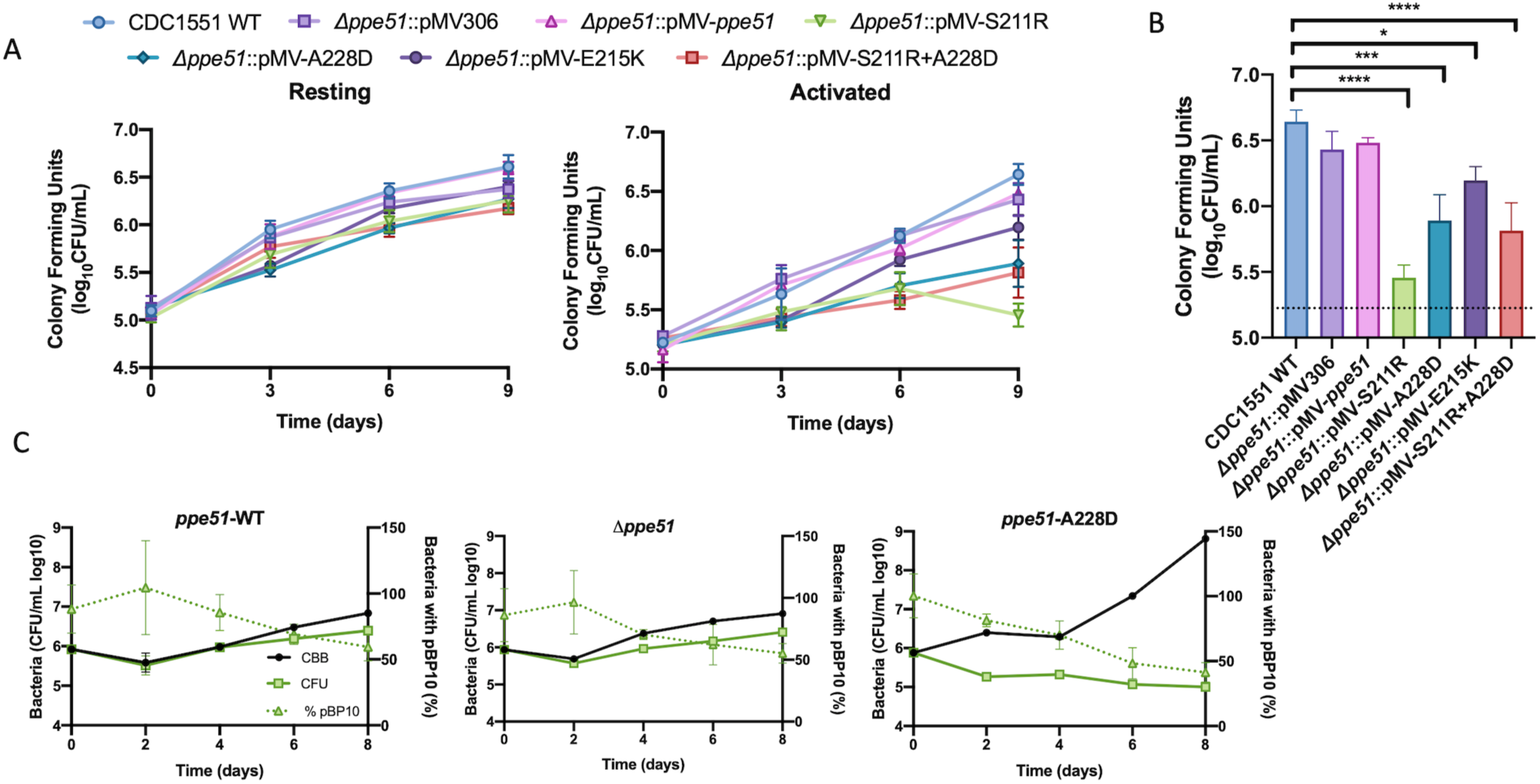
*eag* variants exhibit selectively enhanced replication and reduced survival in activated macrophages. **A)** BMDMs infected with the isogenic panel of CDC1551 *Δppe51* complemented strains and WT CDC1551. Growth is similar for all strains in resting macrophages, but in activated BMDMs, WT, *Δppe51*(empty vector), and *Δppe51*::pMV-*ppe51* exhibit ∼1.25 log increase in growth compared to variant complements which show a lower log increase in growth (∼0.25-1). Data shown was conducted in triplicate and representative of three independent experiments. **B)** Statistical analysis of growth differences between *Δppe51* complemented strains at Day 9 in activated BMDMs. Significance was determined by one-way ANOVA (Tukey’s multiple comparisons test; *P < 0.05, ***P < .001, ****P < 0.0001). **C)** Activated BMDMs infected with native WT CDC1551, *Δppe51*, and A228D variant strains containing the pBP10 replication clock plasmid. CFUs on selective plates were compared to CFUs on nonselective plates and used to calculated frequency of plasmid-bearing bacteria (% pBP10), cumulative bacterial burden (CBB) of total live and dead bacteria, and total enumerated colonies on nonselective plates (CFU).

Rohde *et al*. showed that rapid replication of intracellular Mtb is associated with greater Mtb killing by the macrophage^10^. We observed *in vitro* that variants had enhanced death during replication at acidic pH, and we hypothesized that the *eag* variants may be replicating faster than the WT in macrophages but have lower CFUs due to enhanced death rates. To test this hypothesis, we infected BMDMs with native CDC1551 WT, *Δppe51*, and A228D variant containing the pBP10 plasmid as described previously. Infection was conducted over the course of 8 days with cells lysed and plated for viable CFUs every 2 days. We observed an initial ∼0.5 log decrease in viable CFUs in both WT and *Δppe51* around day 2 that is consistent with observations made by Rohde *et. al*.^59^, and supports their findings that Mtb exhibits delayed adaptation to survive and replicate within macrophages (Figure 6C). Both WT and *Δppe51* then replicated over the course of 8 days inside activated BMDMs as evident by their ∼1 log increase in CFUs starting at day 2. In contrast, the A228D variant lacks this initial adaptation period and instead show a continual ∼1 log decrease in CFUs over the course of 8 days. Calculating the CBB of the A228D variant shows a large difference between the CBB and CFUs, demonstrating that the A228D variant is replicating at a higher rate and dying at an even greater rate. These strains are able to replicate and survive better in resting BMDMs compared to activated BMDMs (Figure S13). We conclude that slowed growth in response to acidic pH inside activated macrophages is necessary for mycobacterial survival and that the *eag* variants do not be sufficiently slow their growth inside macrophages, resulting in enhanced killing. These results also support that the PPE51 variant is promoting uptake of a carbon source during macrophage infection, suggesting that Mtb may metabolize glycerol when growing in macrophages.

### Differential thermal stability of PPE51 and the S211R variant proteins support direct interactions between PPE51 and glycerol

We hypothesized that the *eag* variants promote PPE51 uptake of glycerol by altering PPE51 structure and its affinity for glycerol. Changes in the thermal stability of the protein would provide evidence supporting this hypothesis. C-terminal his-tagged recombinant PPE51 and PPE51 (S211R) variant proteins were expressed and purified from *E. coli* (Figure S14A and B, Table S1). Glycerol was omitted from the reagents used in the purification process and loading dye. In the absence of glycerol, we observed differential stability between the WT and S211R PPE51 variants, with the WT and S211R proteins completely denaturing at 60 °C and 50 °C, respectively, supportive of a significant structural change by the amino acid substitution (Figure 7A and B). In glycerol, the WT protein exhibited enhanced stability, completely denaturing at 65 °C, a shift of 5 °C, and the S211R protein completely denaturing at 60 °C, a shift of 10 °C. These findings support that glycerol/PPE51 interactions, with differential stability shifts dependent on the S211R substitution.

**Figure 7.**
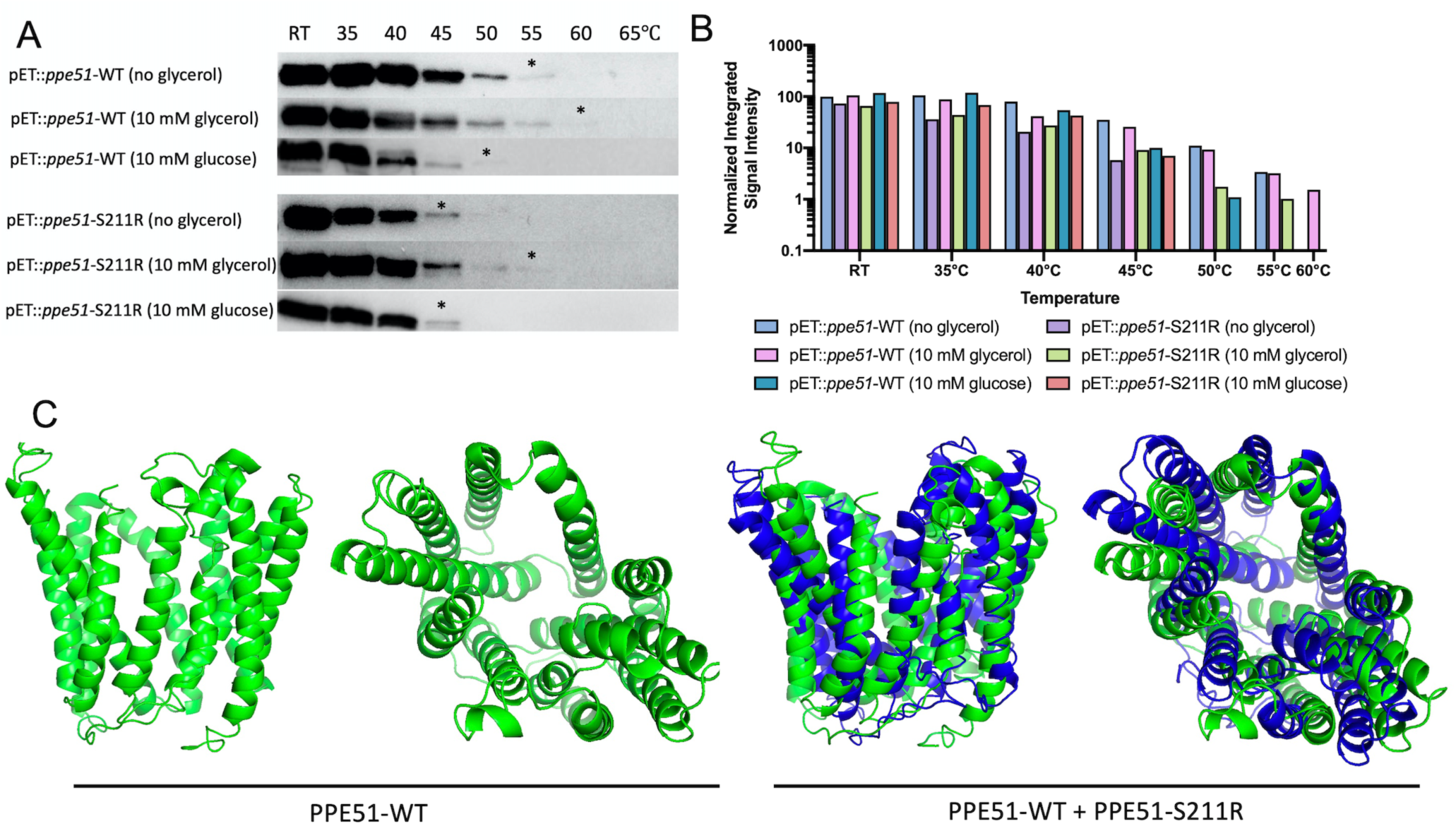
Glycerol differentially interacts with recombinant WT PPE51 or S211R variant proteins. **A)** Recombinant WT PPE51 and S211R proteins were assessed for thermostability under no glycerol, 10 mM glycerol, and 10 mM glucose conditions. The protein was preincubated at room temperature (RT) for 20 minutes and subjected to eight different temperature conditions as indicated for five minutes. Following heating, samples were spun down to pellet the protein precipitate. Soluble protein was removed and analyzed by Western Blotting. * represent the highest temperature where soluble protein was detected. **B)** Signal intensity of individual bands was measured and normalized to the pET::*ppe51*-WT (no glycerol) band at RT. Samples containing glycerol continue to show detectable signal intensity up to 55°C for pET::*ppe51*-S211R and 60°C for pET::*ppe51*-WT. **C.** *in silico* protein structure modeling and function prediction for PPE51. The peptide sequence of PPE51-WT (green) and PPE51-S211R (blue) were analyzed using the Iterative Threading ASSEmbly Refinement (I-TASSER) approach^60^. Both WT and variant PPE51 were modeled without constraint and appear to form a porin-like structure with an inner channel. PPE51-S211R is modeled against the WT to show the slight conformational changes that occur with the introduction of this mutation. PPE51-WT and PPE51-S211R structures received C-scores of -0.86 and -1.24, respectively, which is a measure of structure confidence on a range of -5 (low) to 2 (high)^60^. A model with C-score >-1.5 usually indicates a correct fold.

We previously noted that PPE51 S211R did not promote the growth on glucose and therefore, examined the thermal stability in glucose. We observed reduced stability of the WT protein in glucose, completely denaturing at 55 °C and did not observe any differences in stability with the S211R protein in glucose, supporting the stability shifts are selectively dependent on glycerol (Figure 7A and B). Together, these data show that glycerol selectively increases the thermal stability of PPE51, with enhanced impact on the S211R variant, lending further support for a mechanism whereby PPE51 directly binds glycerol for uptake and acquisition into the Mtb cell.

Based on the *eag* phenotype and differences in thermal stability, we hypothesized that these substitutions may have a significant impact on protein structure and thus conducted *in silico* modeling of PPE51 using the Iterative Threading ASSEmbly Refinement (I-TASSER) server for protein structure and function prediction^60^. The best fit model of PPE51-WT had a moderately high confidence score (C-score) of -0.86 on a scale of -5 (low confidence) to 2 (high confidence) (Figure 7C). Interestingly, threading of the sequence against known transporter structures produced a porin-like model with a possible channel. This model was matched to all structures in the Protein Data Bank (PDB) library. The top 10 proteins from the PDB with the closest structural similarity were all predicted to be nutrient transporter proteins (Figure S14C) An overlay of the S211R variant (blue) with PPE51-WT model shows that the introduction of this substitution confers a noticeable conformation shift in the predicted protein structure (Figure 7C). For PPE51-WT, the top two predicted ligands were maltose and a monoacylglycerol derivative (78N) with one predicted ligand binding site for maltose being within the 18 amino acid residue range of the *eag* variants, located at residue 225 (Figure S14D). Additionally, the predicted top Gene Ontology terms for the molecular function, biological process, and cellular component are hexose:hydrogen symporter activity (GO:0009679), transmembrane transport (GO:0055085), and integral to membrane (GO:0016021), respectively (Figure S14E). While these *in silico* results are predictions, they provide further support for a role with PPE51 acting as a nutrient transporter for Mtb. Furthermore, all *eag* mutations mapped to a single alpha helix on the predicted I-TASSER model, with S211R and E215K located at the top of the predicted channel and A228D located within the center channel structure (Figure S14F) and the substitutions, altered the modeled substrate interaction, further supporting our model for PPE51 variants acting to promote uptake of glycerol by altering the protein structure and ligand interactions.

### PDIM biosynthesis is disrupted in the ppe51 deletion strains

Surprisingly, we found that the *Δppe51* mutant generated in this study does not have the same growth or glycerol uptake compared to previous studies that have generated similar knockouts of knockdowns of *ppe51*^28,41,61^. We also found that the *Δppe51* grew just as well as other strains at pH 7.0 on glycerol (Figures 3B and S8A). This observation was previously made by Wang et. al., who showed that mutations in phthiocerol dimycocerosates (PDIM) biosynthesis were responsible for permeabilizing the mycomembrane and compensating for the loss of functional *ppe51*^28^. We sequenced the genomes of *Δppe51* mutants in both the CDC1551 and Erdman backgrounds and found that both *Δppe51* mutants had evolved mutations in PDIM biosynthesis pathway genes (*ppsC* in *Δppe51*-CDC1551, and *ppsD* in *Δppe51*-Erdman). We confirmed for loss of functional PDIM by radiolabeling it with ^14^C-glycerol and ^14^C-acetate for six days and extracting total lipids for TLC analysis. As expected, we observed loss of functional PDIM accumulation in the *Δppe51*::pMV-EV compared to WT Mtb radiolabeled at both pH 5.7 and pH 7.0 with ^14^C-acetate and ^14^C-glycerol, respectively (Figures 8B and S13). However, despite the occurrence of these PDIM mutations in the *Δppe51* mutants, no PDIM mutations were present in the sequenced *eag* variant mutants used in this study or WT Mtb, highlighting that the *eag* variants remain gain-of-function, dominant mutants (Figure 2A), that selectively promote uptake of glycerol (Figure 2D) and promote enhanced uptake of glycerol and enhanced replication *in vitro* and macrophages (Figures 5A, 3C, 6C, S11B, S12A, S13). Conclusions for the PPE51 knockout or *eag* variants in the *ppe51* knockout background, must take into account that these strains are also PDIM mutants. Overall, we did not observe any differences in the *eag* mutants if they were in a WT or *ppe51*/PDIM mutant background, supporting that these gain-of-function *eag* phenotypes are independent of PDIM levels.

**Figure 8.**
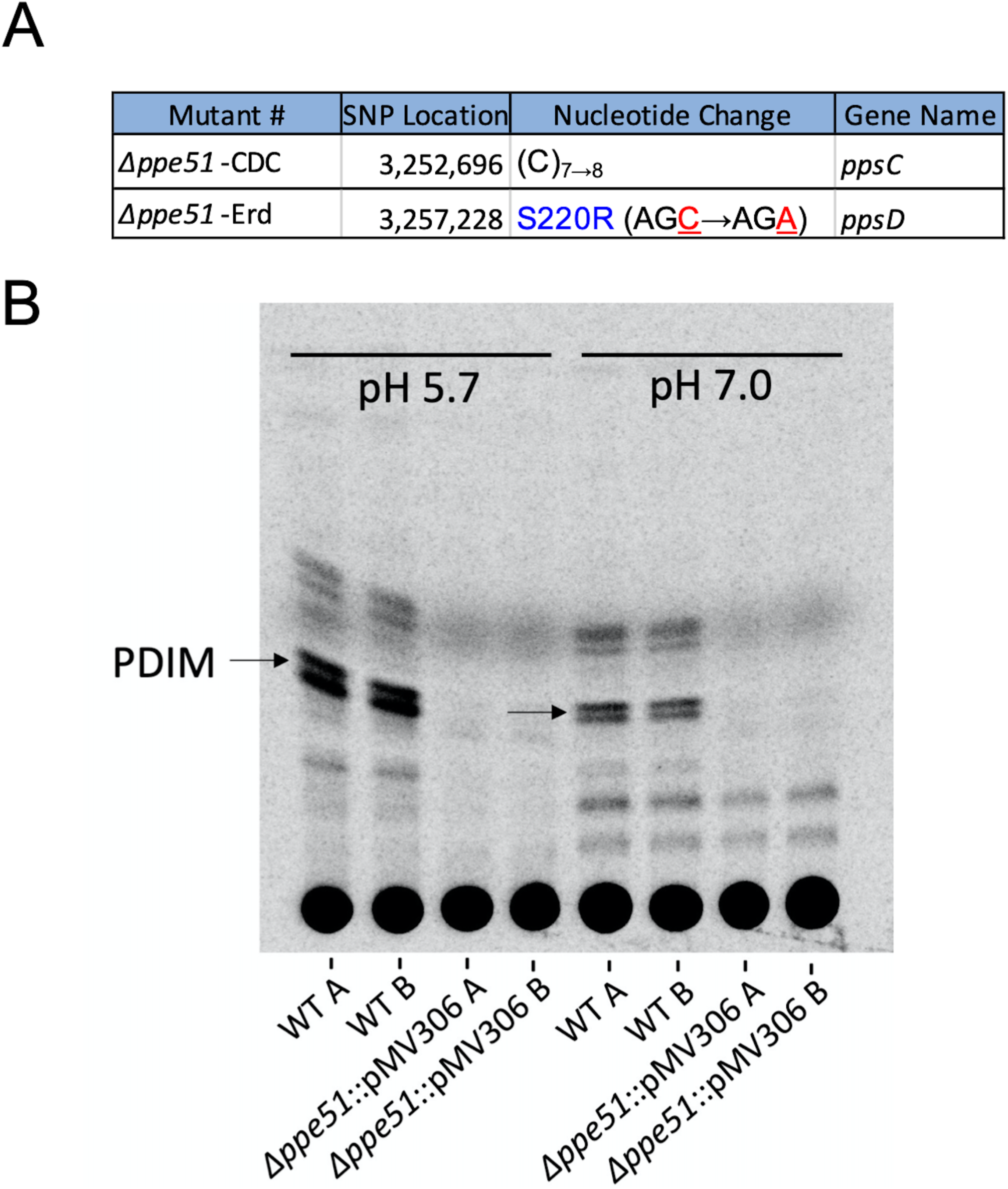
*ppe51* knockout strains contain background mutations that disrupt PDIM biosynthesis. **A)** Whole genome sequencing results of *Δppe51* show a nucleotide insertion in *ppsC* and a point mutation in *ppsD* in CDC1551 and Erdman backgrounds, respectively. **B)** Incorporation of ^14^C-acetate into PDIM at acidic and neutral pH. PDIM is indicated with an arrow and accumulates in the WT strain at both pH 5.7 and pH 7.0 but is absent in the *ppe51* knockout mutant. Strains were analyzed in duplicate with representative results being shown.

## Discussion

Mtb exhibits complex regulatory and physiological adaptations when grown in acidic environments, including changes in growth rate. The underlying basis of slowed growth in mildly acidic environments is still not fully resolved, but appears to be associated with metabolic and redox stress, that may be linked to balancing cytoplasmic pH-homeostasis and respiration^11^. Providing specific carbon sources, such as pyruvate or acetyl-CoA, relieve this metabolic stress and enable Mtb to grow similarly well at acidic and neutral pH^16,18^. However, it has been puzzling as to why Mtb cannot grow on glycerol at acidic pH, as it has a carbon source and oxygen, everything it needs to grow. In this study, we found that Mtb limits uptake of glycerol at acidic pH to restrict its growth and that mutations in *ppe51* promote uptake of glycerol at acidic pH and enable growth. That is, Mtb can grow well at acidic pH on glycerol, but has adapted instead to stop growth.

We further show that this pH-dependent metabolic adaptation is required for pathogenesis. Selectively in activated macrophages, where the pH of the phagosome is more acidic, we observed a virulence defect in strains expressing the *eag* variants. Notably, using a replication clock plasmid, we found that *eag* variants have enhanced growth in macrophages, but even greater killing, the balance of which results in reduced fitness. Thus, slowed growth in macrophages, in an activation dependent manner is dependent on the restriction of metabolism at acidic pH, and that PPE51 variants overcome this restriction to the detriment of the pathogen. This finding supports that the nutrient imported by the PPE51 variant is relevant to the macrophage environment. We showed that the variants specifically promote uptake of glycerol, therefore, it is plausible that glycerol is a key regulator of Mtb growth in the macrophage. It has been previously shown that Mtb can uptake TAG in macrophages^62^, TAG is abundant in granulomas^63^, and Mtb exports the TAG lipase LipY^64^, therefore, it is possible glycerol is released from TAG during infection, and restriction of glycerol uptake plays an important role in slowing growth during infection. Studies examining the interactions of PPE51 *eag* variants, LipY and glycerol metabolism genes during pathogenesis will be undertaken to test this hypothetical model.

It is a striking finding that all of the *eag* mutants selected were in *ppe51* and that they all clustered with a highly conserved region of 18 amino acid residues (residues 211-228). Three single amino acid substitutions (S211R, A228D, and E215K) greatly altered WT *ppe51* function and promoted growth under acid stress when given the non-permissible carbon source, glycerol. S211R was able to confer the greatest enhanced growth, whereas A228D conferred moderate enhanced growth and E215K exhibited the least amount of enhanced growth, comparatively (Figure 2B). The growth phenotypes of the native mutant alleles were further recapitulated in overexpression studies in a WT Mtb background as well as a *Δppe51* background, where again we observed overall greater *eag* with the S211R variant compared to A228D and E215K (Figures 2A, 3A, S8A). Given that the phenotype was conserved in PDIM containing strains (the initially isolated mutants and the overexpressors in the WT) and PDIM lacking strains (the *ppe51* deletion mutants), this demonstrates that the gain-of-function phenotype is independent of PDIM. This region of PPE51 may play a key role in protein-substrate interactions, and indeed with recombinant proteins, we observed differential stability in the variant protein and its interaction with glycerol. Interestingly, the structural modeling showed substitutions in this region altered the predicted ligand of the modeled transporter, supporting further study of this critical region for modulating PPE51-ligand interactions.

Another key finding of this study is that glycerol uptake is restricted at acidic pH. Data supporting this conclusion include the reduced uptake of radiolabeled glycerol at acidic pH as compared to neutral pH (Figures 5A and S12AB), the dependence of glycerol concentration and pH in regulating growth (Figures 4, S11A and S11B), and the ability of PPE51 variants to enhance growth and glycerol uptake at acidic pH (Figures 5A and S12AB). How Mtb restricts glycerol uptake is still not known, but it is puzzling that PPE51 is strongly induced at acidic pH and counter to a model where PPE51 promotes in glycerol uptake, but Mtb restricts glycerol uptake at acidic pH. This contradiction remains unresolved and points to a new known unknown of Mtb metabolism restriction at acidic pH. Notably, growth on glycerol-containing mixtures can exceed growth compared to growth on glycerol alone^65^, suggesting that Mtb may need to restrict glycerol to regulate its growth while consuming other carbon sources it encounters during infection.

We identified that the *eag* variants selectively enabled growth on glycerol alone compared to WT Mtb (Figure 2D). The identification of this carbon specificity with PPE51 *eag* variants implies a putative role for PPE proteins in nutrient acquisition, a model that is strongly supported by data put forth by Ates *et. al.*, Mitra *et. al*. and Wang *et al*. These studies showed that PE and PPE proteins located at the cell envelope and cell surface play a vital role in nutrient uptake for Mtb. Ates *et. al.* provides strong evidence that the type VII secretion system, ESX-5, is essential for mycobacterial growth and nutrient uptake. In this study, essentiality of ESX-5 could be rescued by altering cell wall lipid composition or introducing the *M. smegmatis* outer membrane porin, *mspA*, which mediates cell wall permeability and influx of hydrophilic nutrients^66–68^. ESX-5 mutations in *M. marinum* result in significantly reduced growth on medium with Tween-40 or Tween-80 as the sole carbon source, and the ESX-5 mutant strain exhibits significantly impaired uptake of fluorescently-labeled fatty acids compared to WT and complemented strains^34^. These data support ESX-5 facilitating the uptake of fatty acids to be used as a carbon source through the secretion of PE and PPE proteins. In support of Ates’ ESX-5 substrate nutrient influx hypothesis, Mitra and colleagues^26^ showed direct evidence tying PE and PPE proteins to iron acquisition. Mitra identified Mtb transposon mutants that were resistant to a toxic heme analog^26^. The mutants were in three previously uncharacterized genes of which two were PPE proteins, PPE36 and PPE62^26^. Furthermore PPE62 was shown to be surface-accessible and predicted structure indicates that it may form a β-barrel that resembles *Haemophilus influenzae* heme cell surface receptor^26,69^ and that heme transport is facilitated into the cell by the periplasmic lipoprotein DppA^70^. Finally, the Wang *et al.* study provided direct evidence that PPE51 is exported to the mycomembrane to promote uptake of glycerol and glucose, possibly by acting like a porin. Notably, loss of function *ppe51* mutants have altered sensitivity to antibiotics, including pyrazinamide^71^ and meropenem^72^, suggesting that PPE51 mediated impacts on carbon source uptake or mycomembrane permeability play a role in drug susceptibility, supporting further studies of PPE51 as a target for potentiating antibiotics.

Here, we present a model that integrates the current understanding of PE and PPE nutrient acquisition with our findings (Figure 9), wherein PPE51 embeds itself into the outermost layer of the cell envelope and is surface accessible to glycerol^28,42,73^. Gene expression profiling data supports induction of *ppe51* by *phoP* and acidic pH^15,18^. Phylogenetic evidence shows that PPE51 is duplicated alongside ESX-5^32^, which has been shown to mediate the secretion of most PE/PPE proteins in *M. marinum*, including PPE51^34,38^. We propose that an unknown periplasmic nutrient transporter helps mediate the import of glycerol across the plasma membrane and into the cell from initial import by PPE51. *pe/ppe* families have high variation rates between *Mycobacterium tuberculosis* complex (MTBC) genomes with *ppe51* being the single exception in showing almost no variation^74^. However, under the selective pressure of our screen (Figure 1A), we have shown that we can select for mutations that enhance PPE51’s proposed uptake of glycerol (Figure 9). Furthermore, our initial *in silico* modeling of PPE51 suggests that it can form a porin-like structure consistent with a role in transport and ligand-binding sites for carbon nutrient sources (Figure 8C). Based on these data, we further propose an *eag* mutant model, whereby the *eag* amino acid substitutions introduce conformational changes that allow for a possible PPE51-porin structure to widen or enhance the binding the glycerol, allowing enhanced transport through the mycomembrane.

**Figure 9.**
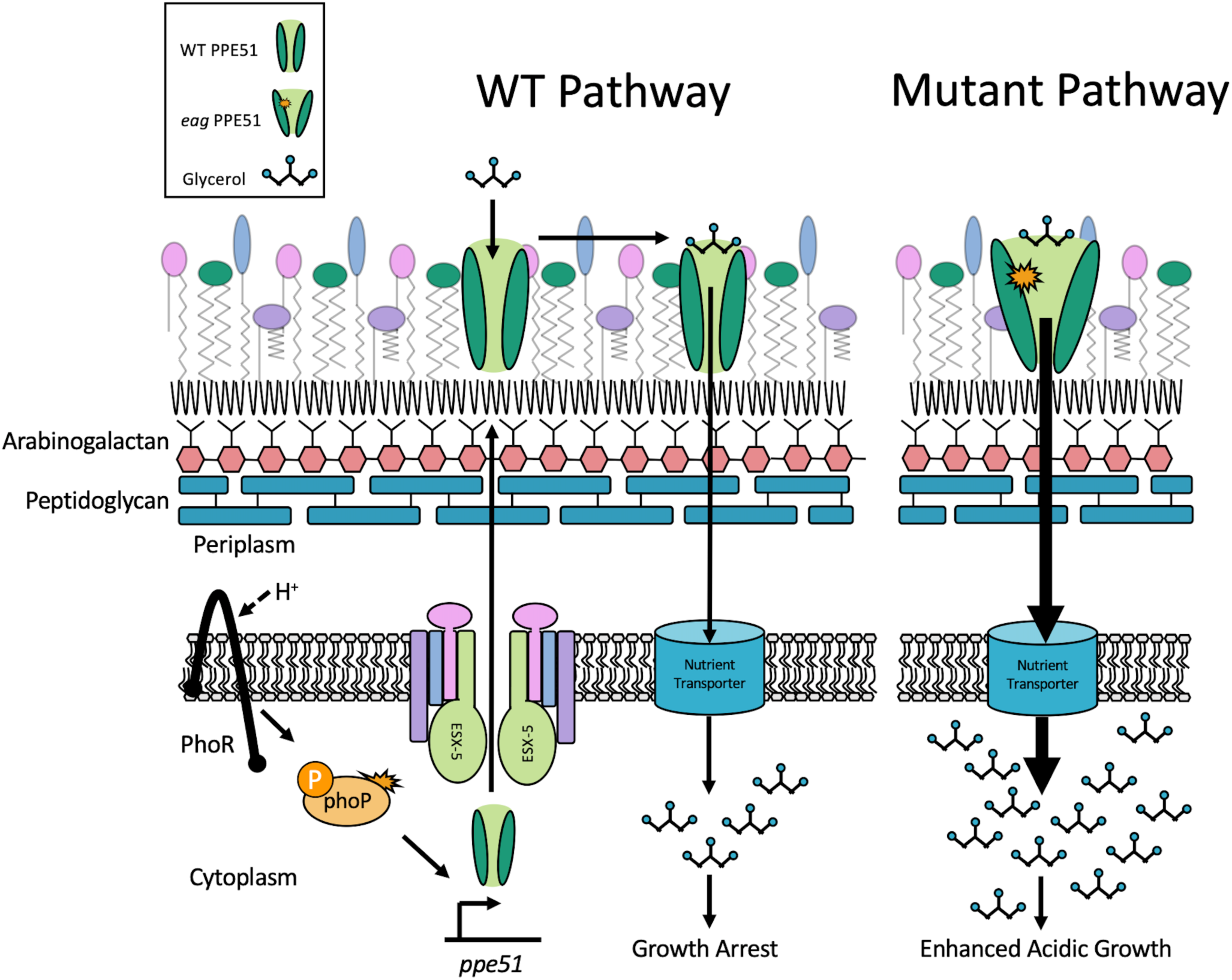
A proposed model for the role of *ppe51* and *eag* variants in glycerol acquisition. Presented is a hypothetical model, in which *ppe51* expression is induced by PhoP under acidic conditions. PPE51 is thought to be secreted through ESX-5 and embeds itself into the mycomembrane, making itself surface-accessible. At this interface, it could interact with glycerol and promote transport across the mycomembrane (WT Pathway). PPE51 variants may function by having an altered channel opening or ligand binding surface, allowing for enhanced glycerol transport across the mycomembrane and leading to the enhanced growth phenotype observed during acid growth arrest (Mutant Pathway).

This study has focused on the role of the *eag* PPE51 variants, and the not the Δ*ppe51* mutant, due to confounding mutations in PDIM in the deletion strains. It is interesting that both deletion mutants (in Erdman and CDC1551) evolved these mutations during the construction of the mutants and suggests there may have been a selective advantage for the mutations. Indeed, Wang et al., showed that *ppe51* knockouts only had a glycerol uptake phenotype when the PDIM was restored in the mutant. This finding is consistent with our observation that the Δ*ppe51* mutants in this study did not have a growth defect in glycerol, presumably due to the lack of PDIM, whereas the *ppe51* mutants in the Wang *et al.,* study were defective for growth. Given the conservation of the *eag* mutants in strains with or without PDIM, we conclude that PDIM level do not appreciably impact the enhanced uptake of glycerol in *eag* variants. However, it is also possible that differences for the PPE51 mutants between this study and the others may be driven by genomic differences. Both Wang *et. al* and Korycka-Machała *et. al*. used the H37Rv Mtb strain for their knockout and CRISPRi knockdown studies, respectively. However, sequence analysis of the region directly upstream of *ppe51* in both CDC1551 and Erdman compared to H37Rv shows an almost total deletion of the *ppe50* gene preceding *ppe51*. The *ppe50* sequence is also not present anywhere else in the CDC1551 or Erdman genome except for a matching 66 bp sequence that precedes *ppe51* in both genomes. The large sequence difference in the *ppe51* promoter region between strains could imply an additional reason why we see strong phenotypic growth differences between our respective growth screens of *ppe51* knockouts.

## Acknowledgements

We thank members of the Abramovitch lab for critical reading of the manuscript and Prof. David Sherman for sharing the pBP10 clock plasmid. This research was supported by a grant from the NIH-NIAID (R01AI116605) and AgBioResearch.

## Author Contributions

S.J.D., J.J.B., and R.B.A. conceived the project. S.J.D performed all of the experimental studies.M.M. conducted thermal stability assay studies. S.J.D. and R.B.A. wrote the manuscript. All authors reviewed the manuscript.

---

**Supplemental Figure 1.**
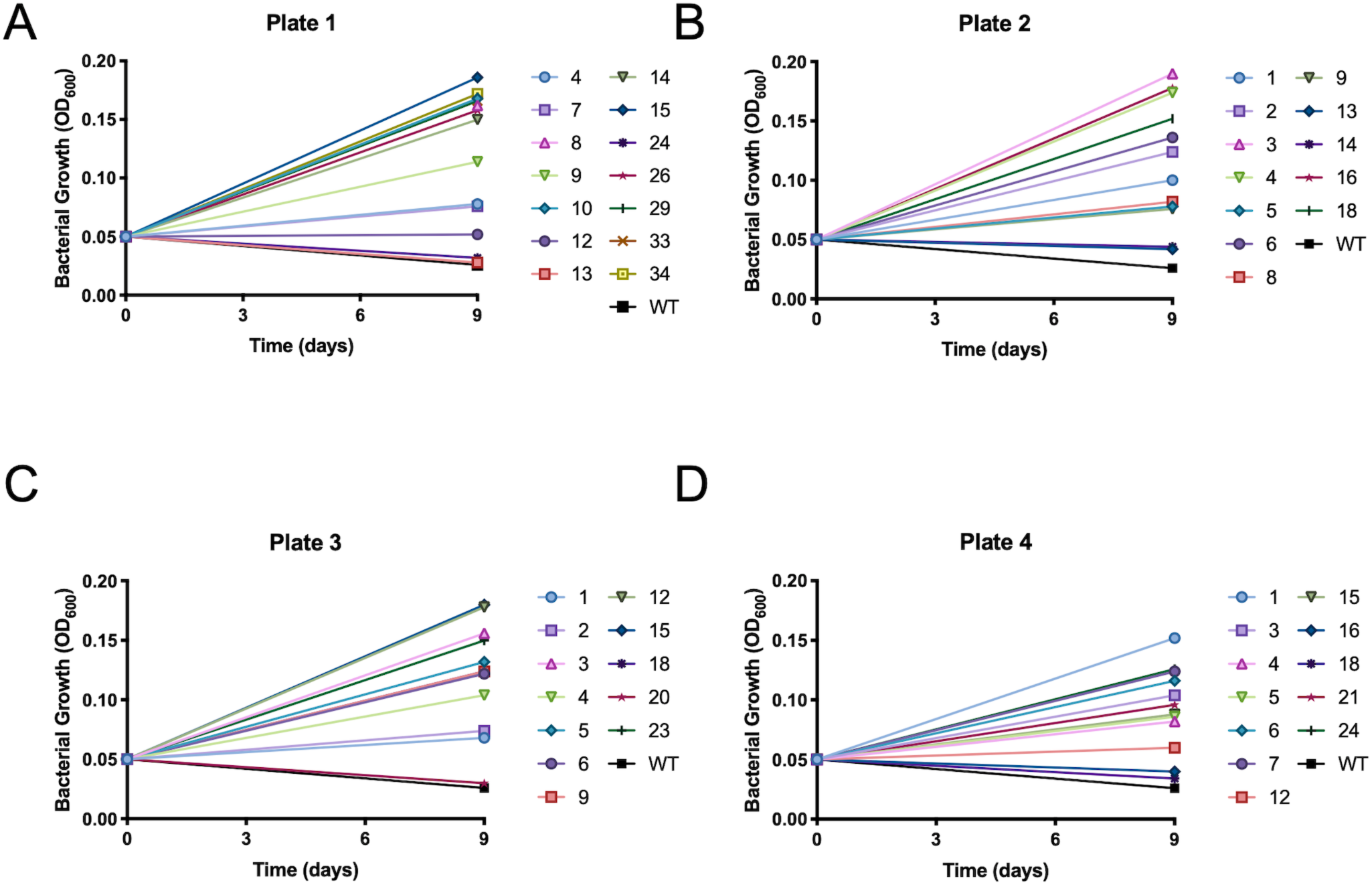
Enhanced acid growth confirmation of mutants isolated from WT Erdman genetic screen. Single colony isolates from Plate 1 (A), Plate 2 (B), Plate 3 (C), and Plate 4 (D) were grown in liquid MMAT (pH 5.7) with glycerol and compared to the WT for the enhanced acid growth phenotype. Each symbol represents the numbered colony isolated from the acid growth arrest plates.

**Supplemental Figure 2.**
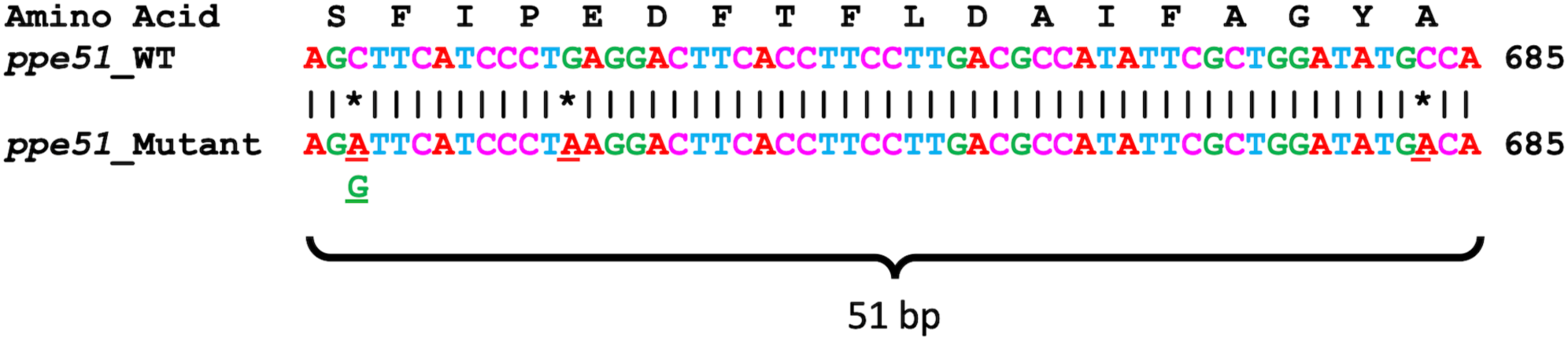
SNP sites in ppe51. SNP mapping identified three separate mutations within a 50 bp region of ppe51. Underlined and starred bases represent the SNP position (bp). S211R substitution had two SNPs as denoted by the guanine (G) underneath.

**Supplemental Figure 3.**
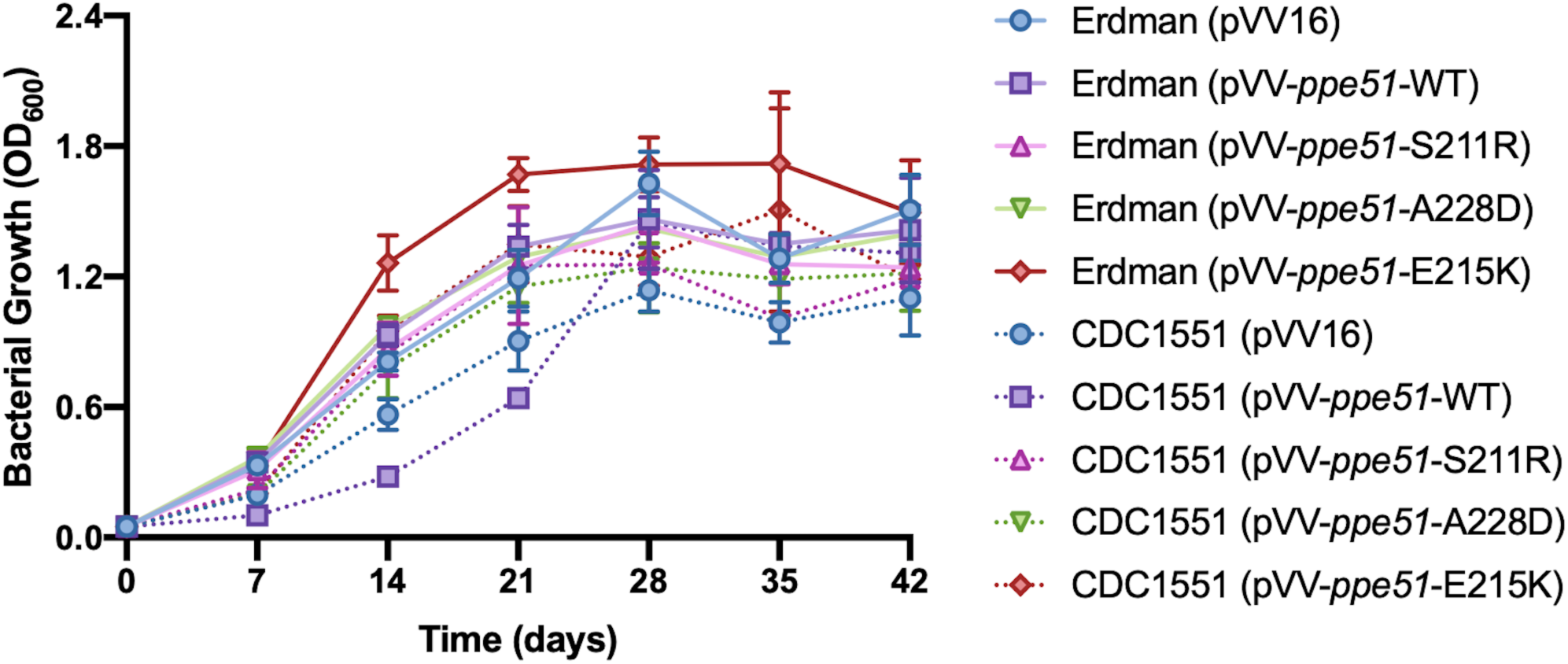
Growth curve of pVV16 overexpression constructs (CDC1551 and Erdman) in minimal media at pH 7.0 with 10 mM glycerol.

**Supplemental Figure 4.**
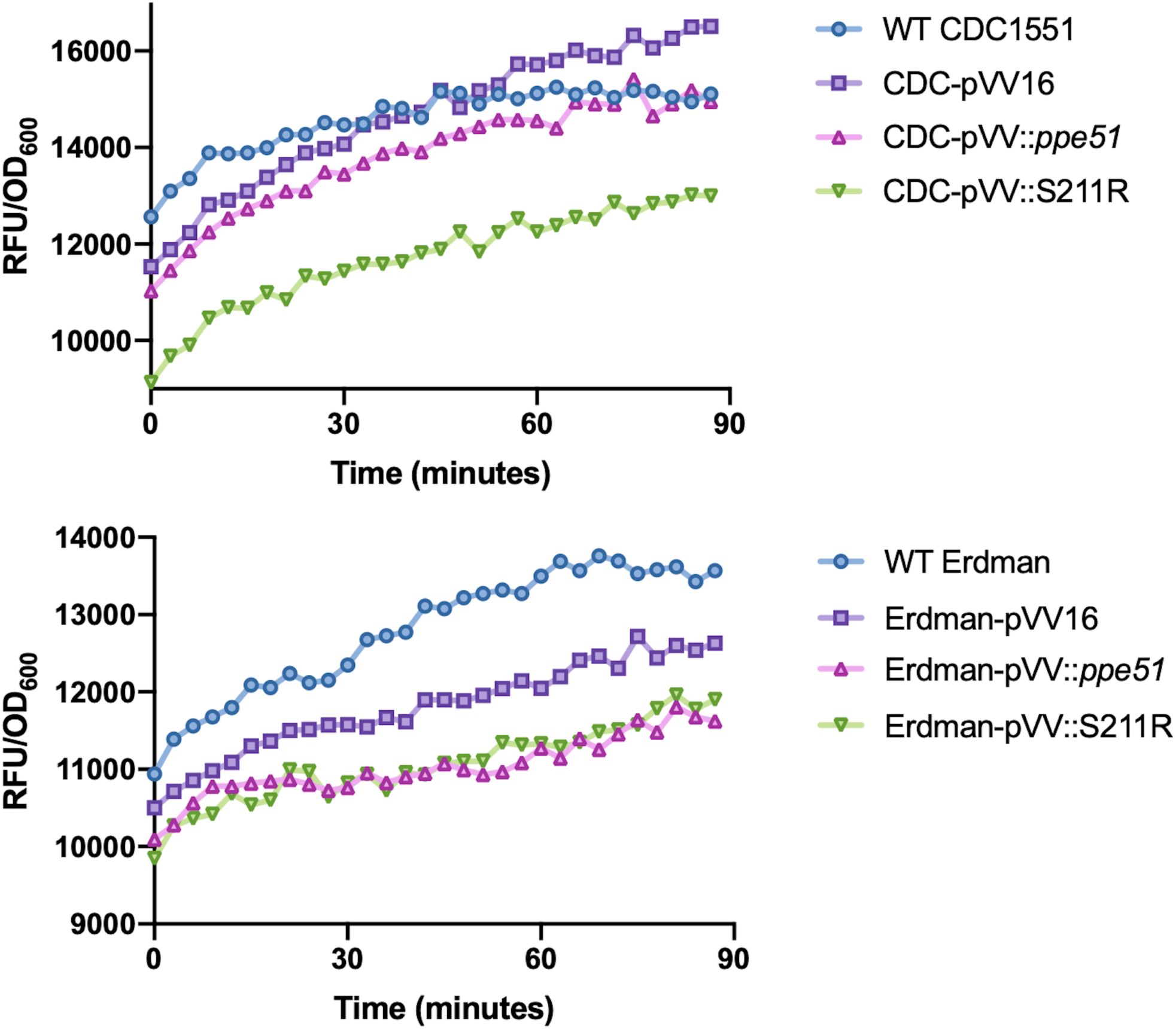
Accumulation of EtBr by Mtb and pVV16 overexpression constructs. pVV16 overexpression constructs (CDC1551 and Erdman) were incubated with Ethidium bromide (EtBr) for 90 minutes. EtBr fluorescence was measured every 3 minutes using the excitation wavelength (530 nm) and emission wavelength (590 nm). Samples were measured in triplicate.

**Supplemental Figure 5.**
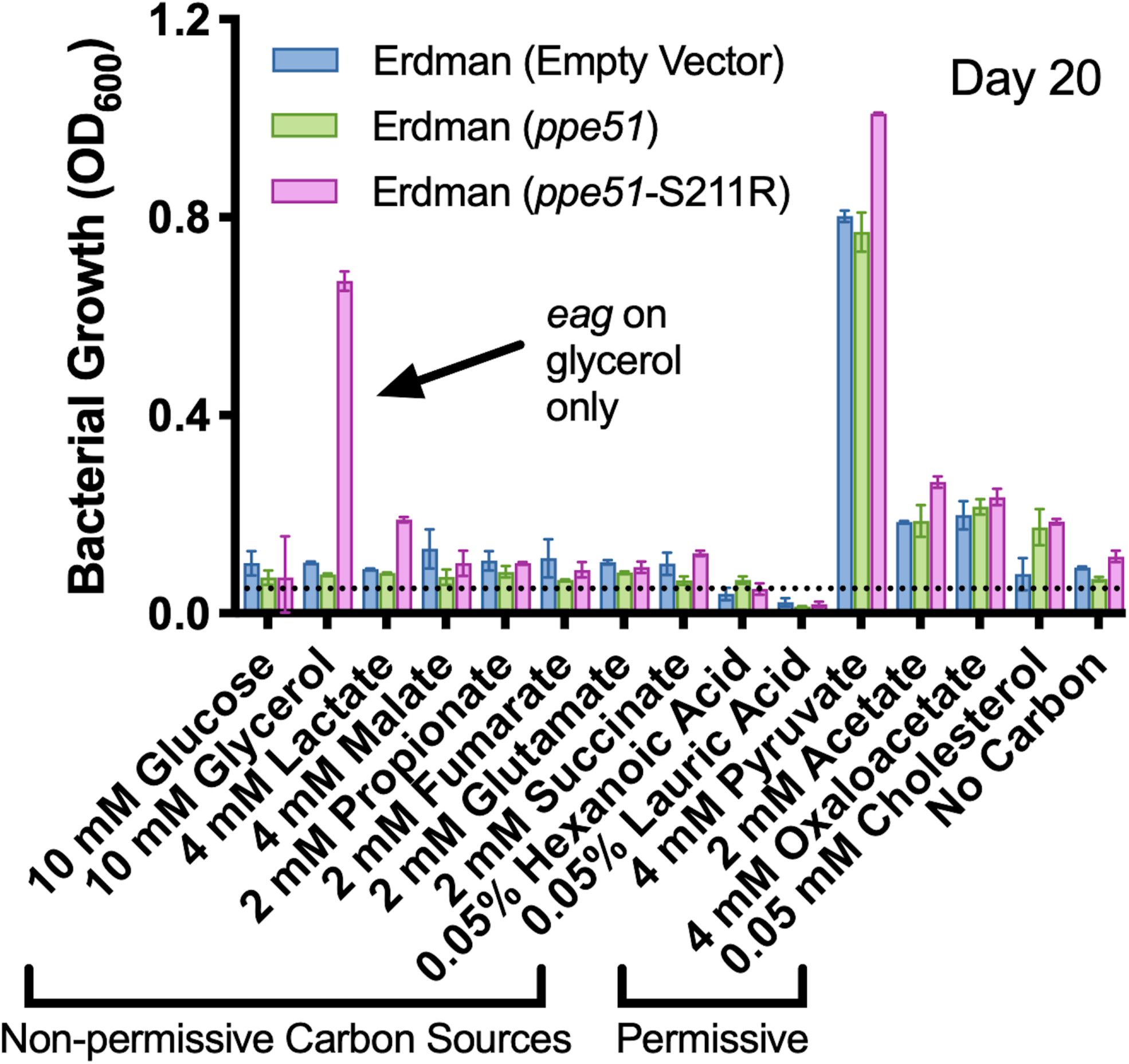
Analysis of the S211R variant growth on various carbon sources. Erdman overexpression strains were grown in MMAT medium (pH 5.7) in the presence of various growth-permissive (i.e. pyruvate acetate, OA) and non-permissive (i.e. glucose, propionate, lactate) carbon sources. ppe51-S211R (pink bars) growth is carbon source specific and only exhibits enhanced growth on glycerol, a normally non-permissive carbon source at pH 5.7. Growth on permissive carbon sources is not impacted by ppe51-S211R. The horizontal dotted line indicates the starting density of 0.05 OD600.

**Supplemental Figure 6A.**
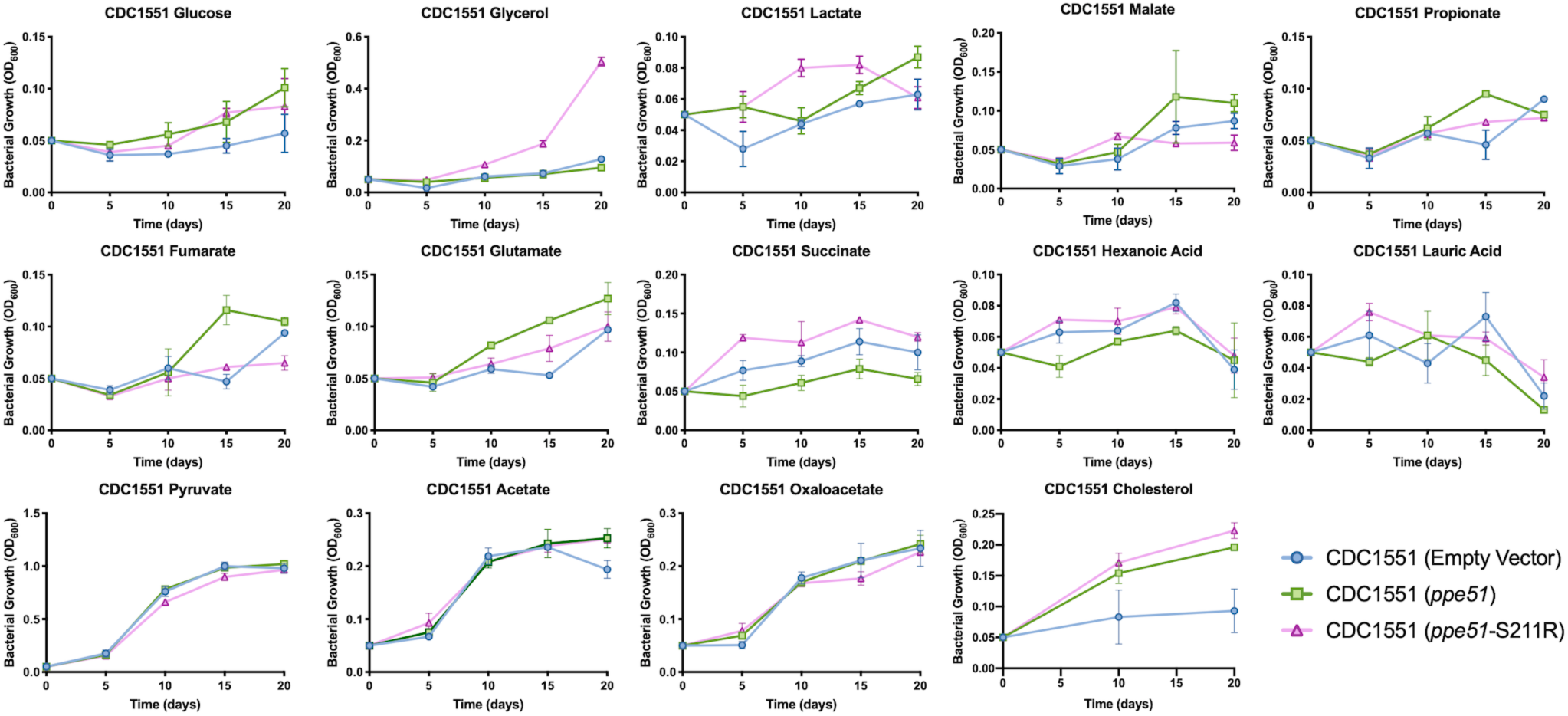
Growth curves of CDC1551 overexpression strains on individual carbon sources. Each strain was grown in duplicate for 20 days on their respective carbon source in minimal media buffered to pH 5.7.

**Supplemental Figure 6B.**
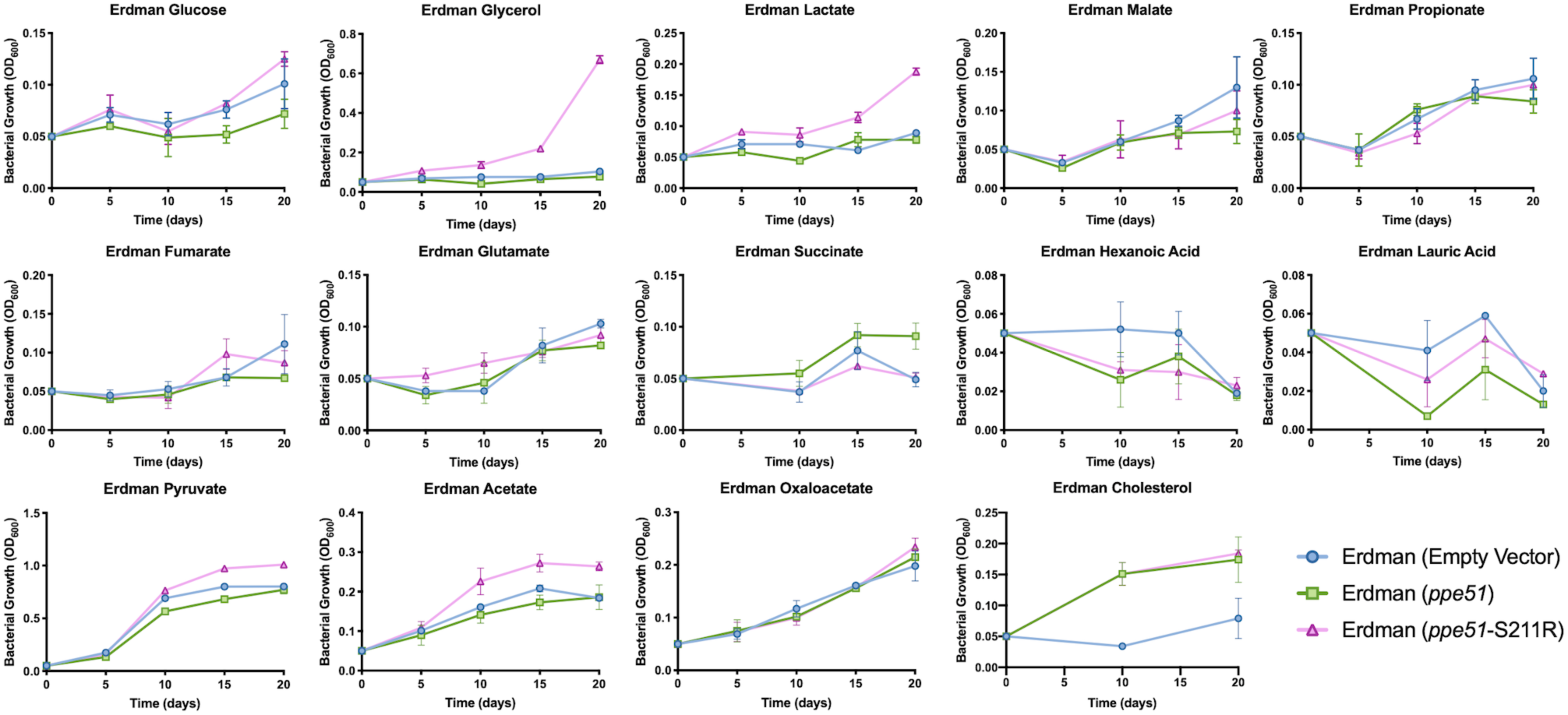
Growth curves of Erdman overexpression strains on individual carbon sources. Each strain was grown in duplicate for 20 days on their respective carbon source in minimal media buffered to pH 5.7.

**Supplemental Figure 7.**
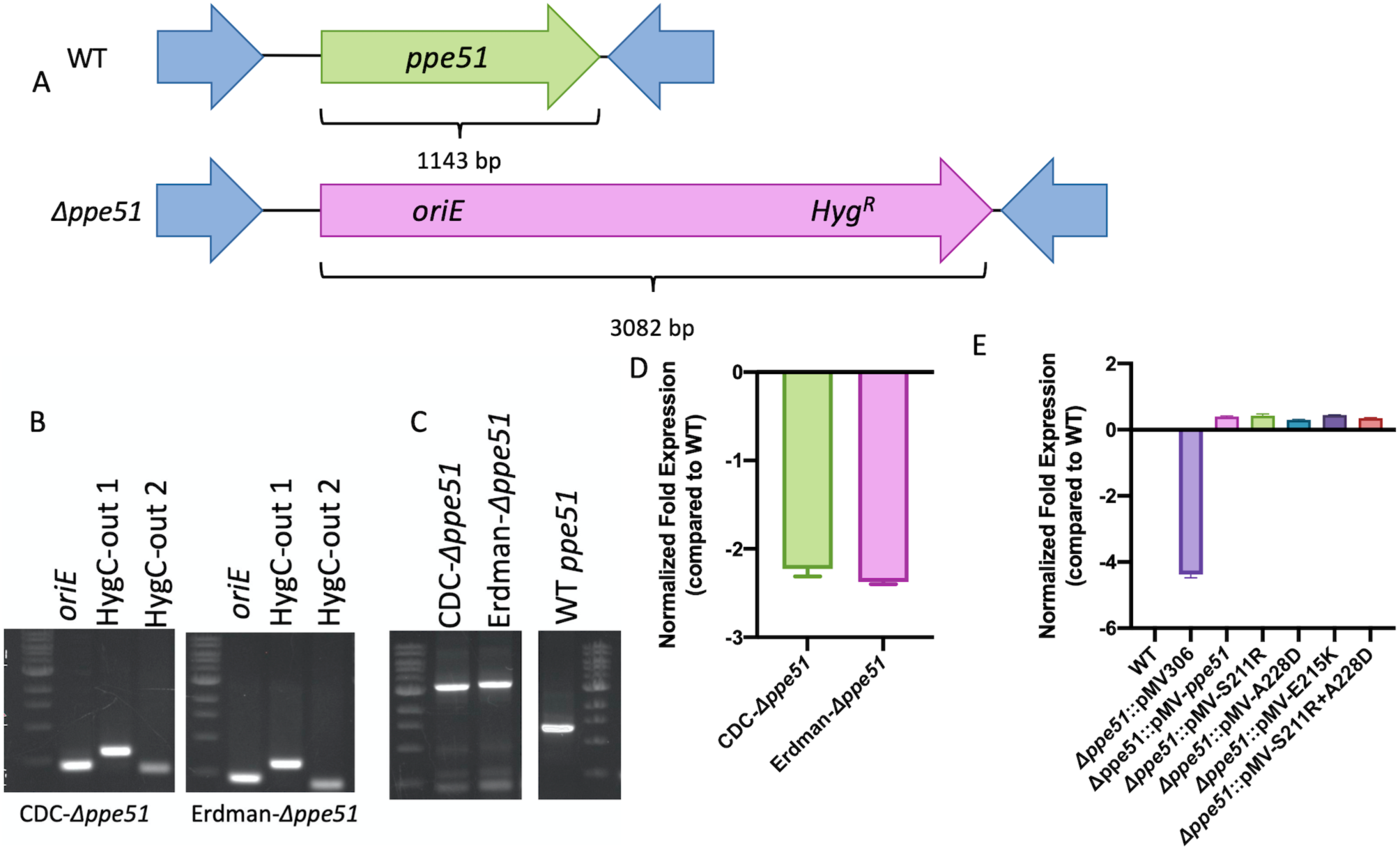
Construction of ppe51 deletion mutant in Mtb CDC1551 and Erdman. A) Schematic of chromosomal ppe51 and the subsequent ORBIT48-promoted deletion of the ppe51 target gene, attP replacement, and plasmid integration containing hygromycin resistance for selection. B) PCR amplification of the 5’(oriE) and 3’ (HygC-out1/2) junctions of CDC1551 and Erdman Δppe51. Positive bands were confirmed by sanger sequencing. C) PCR analysis of the integration site of the payload plasmid (pKM464). pKM464 is 3082 bp which is consistent with the size of the bands observed in the ppe51 deletion mutants compared to WT ppe51 which is 1143 bp. D) qRT-PCR analysis confirming the ppe51 knockout in CDC1551 and Erdman. Total RNA was collected after samples were grown for six days in minimal media buffered to pH 5.7. Fold expression was normalized to the respective WT strains. Error bars represent the standard deviation of three technical replicates. Deletion mutants typically exhibited a non-specific primed Ct ∼30 cycles compared to WT which had a Ct of ∼20 cycles. E) qRT-PCR analysis confirming the presence of the ppe51 gene in complemented Δppe51. Fold expression was normalized to WT Mtb. Error bars represent the standard deviation of three technical replicates. Complemented strains had Ct ∼14 cycles compared to Δppe51 which had non-specific primed Ct ∼30 cycles.

**Supplemental Figure 8.**
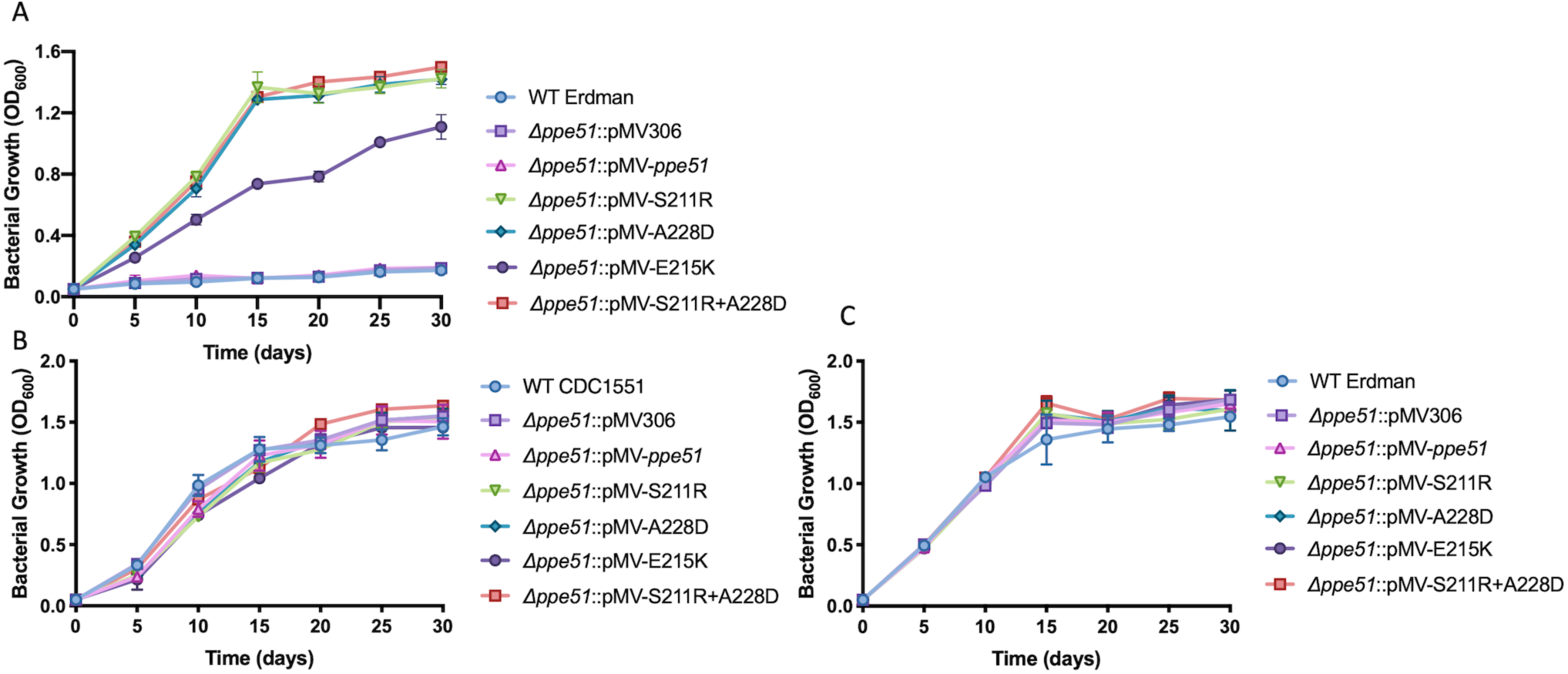
Growth of complemented Δppe51 strains in A) minimal media at pH 5.7 with 10 mM glycerol (Erdman), B) minimal media at pH 7.0 with 10 mM glycerol (CDC1551) and C) minimal media at pH 7.0 with 10 mM glycerol (Erdman). Error bars represent standard deviation of three technical replicates.

**Supplemental Figure 9.**
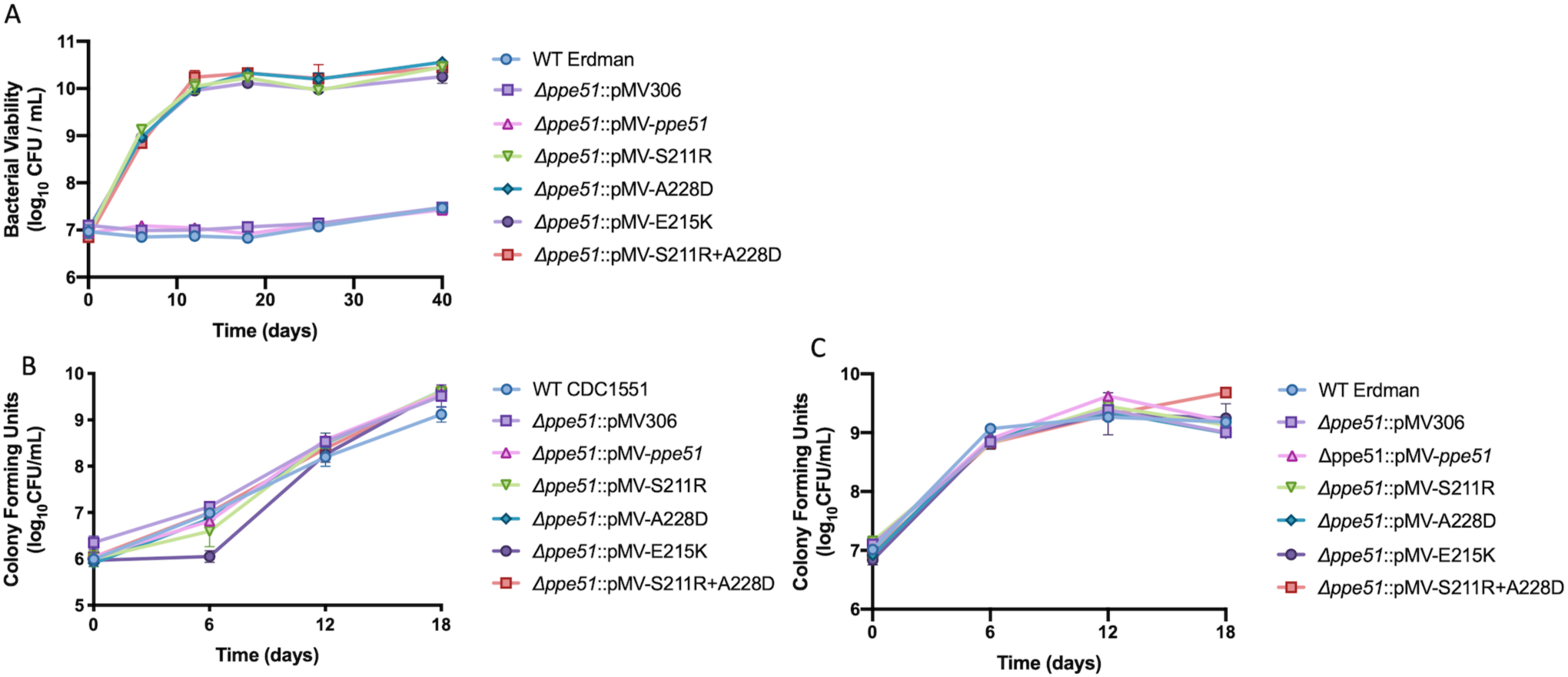
Viability of complemented Δppe51 strains in A) minimal media at pH 5.7 with 10 mM glycerol (Erdman), B) minimal media at pH 7.0 with 10 mM glycerol (CDC1551) and C) minimal media at pH 7.0 with 10 mM glycerol (Erdman). Error bars represent standard deviation of three technical replicates.

**Supplemental Figure 10.**
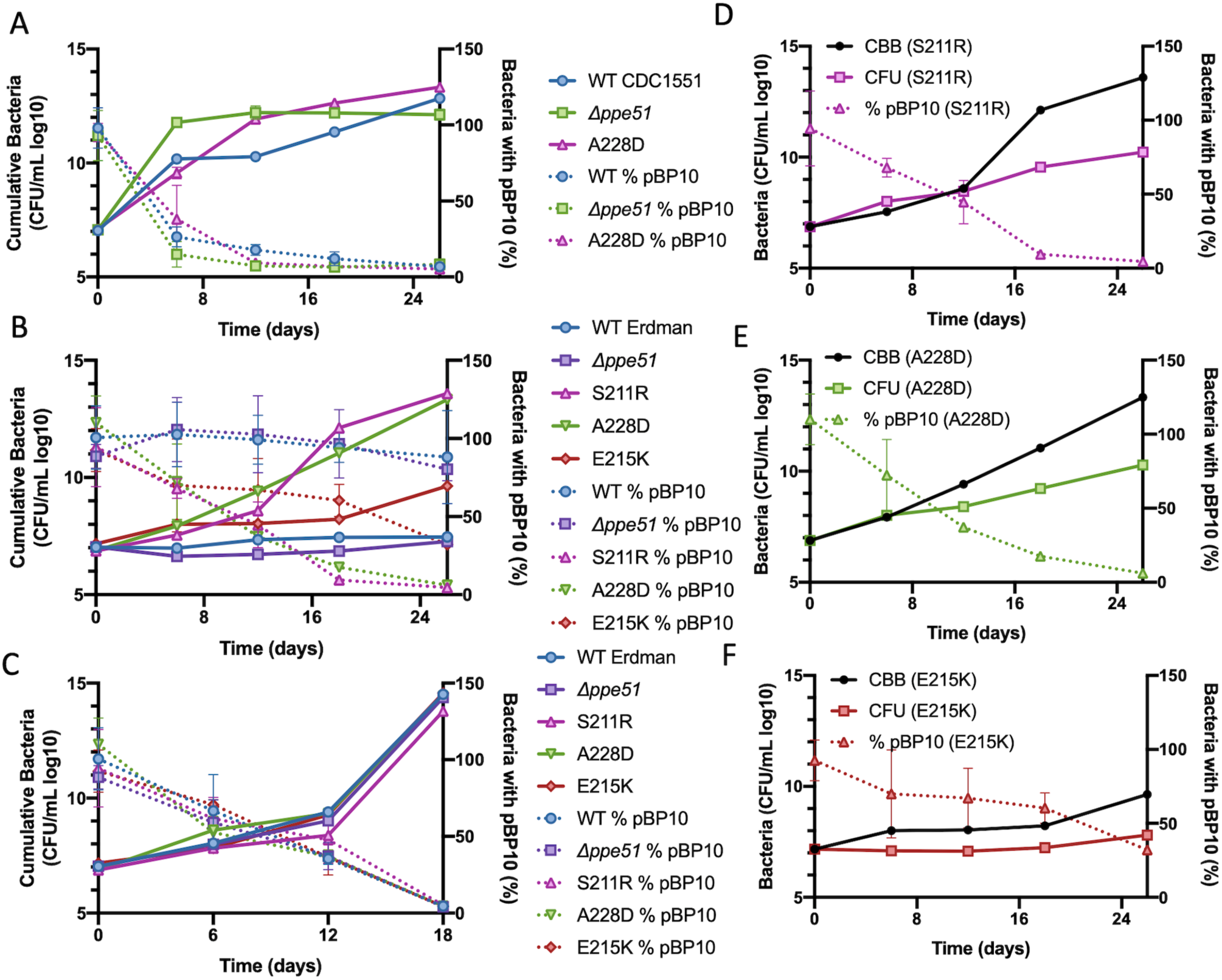
In vitro replication dynamics of CDC1551 eag variants (pH 7.0) and Erdman eag variants (pH 5.7 and pH 7.0). A) All CDC1551 Mtb strains, WT, Δppe51, and the native variant allele, A228D, continues to replicate at neutral pH in minimal media. To estimate replication dynamics of the indicated strains, plasmid frequency data was obtained from CFU counts (right axis, dotted lines). CFUs of plasmid-free and plasmid-bearing strains were then used to calculate cumulative bacterial burden (CBB) of total live, dead, or degraded Mtb (left axis, solid lines). B) Erdman strains containin the native variant alleles, S211R, A228D, and E215K continue to replicate at pH 5.7 in minimal media, albeit E215K exhibits a more reduced capacity for replication compared to S211R and A228D. WT and Δppe51 cease replication. Plasmid frequency (right axis, dotted lines) and cumulative bacterial burden (CBB) (left axis, solid lines) are shown. C) All Erdman Mtb strains, WT, Δppe51, and the native variant alleles (S211R, A228D, and E215K), continue to replicate at neutral pH in minimal media. Plasmid frequency (right axis, dotted lines) and cumulative bacterial burden (CBB) (left axis, solid lines) are shown. Replication dynamics of the native Erdman D) S211R variant, E) A228D variant, and F) E215K variant (comparing CBB (cumulative bacterial burden), CFU (total CFUs from nonselective plating), and % pBP10 (percentage of bacteria carrying plasmid).

**Supplemental Figure 11A.**
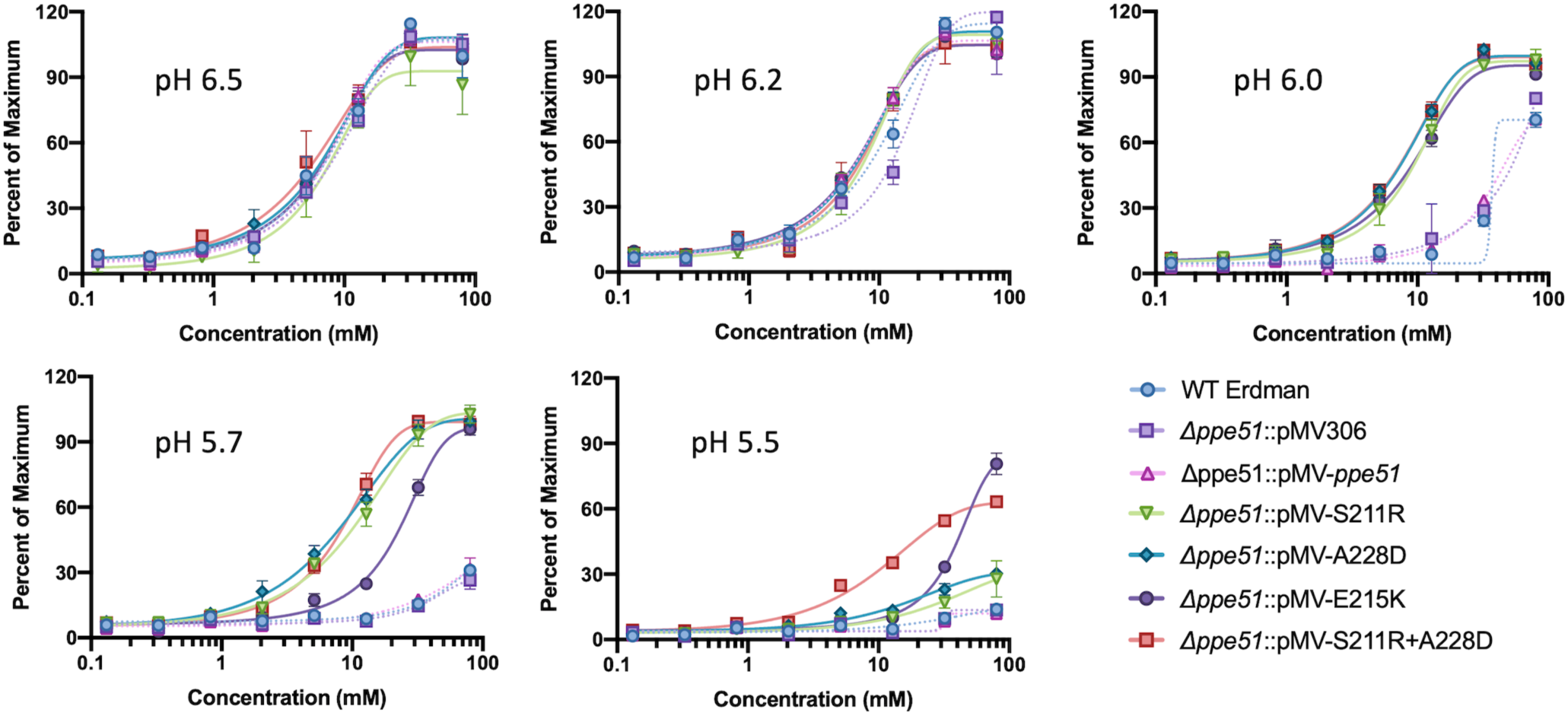
Mtb shows growth restriction at low pH in Erdman. Growth of WT Erdman, Δppe51 (empty vector), and Δppe51 complemented strains in minimal media supplemented in a dose-dependent manner with glycerol and buffered to one of five pH levels (pH 6.5, 6.2, 6.0, 5.7, or 5.5). All strains exhibit a reduced capacity for growth starting ∼2 mM glycerol compared to higher glycerol concentrations. At decreasing pH, WT, Δppe51(empty vector), and Δppe51::pMV-WT restrict their ability to uptake glycerol, whereas any variant complement is able to maintain glycerol uptake. However, restricted growth can be rescued at high concentrations of glycerol (∼80 mM) at pH 5.7 for WT, Δppe51(empty vector), and Δppe51::pMV-ppe51, and pH 5.5 for variant complements. Growth analyses were performed at Day 14 following initial inoculation with data being shown as percent of the maximum well-growth. All conditions were conducted in triplicate and representative of multiple independent experiments.

**Supplemental Figure 11B.**
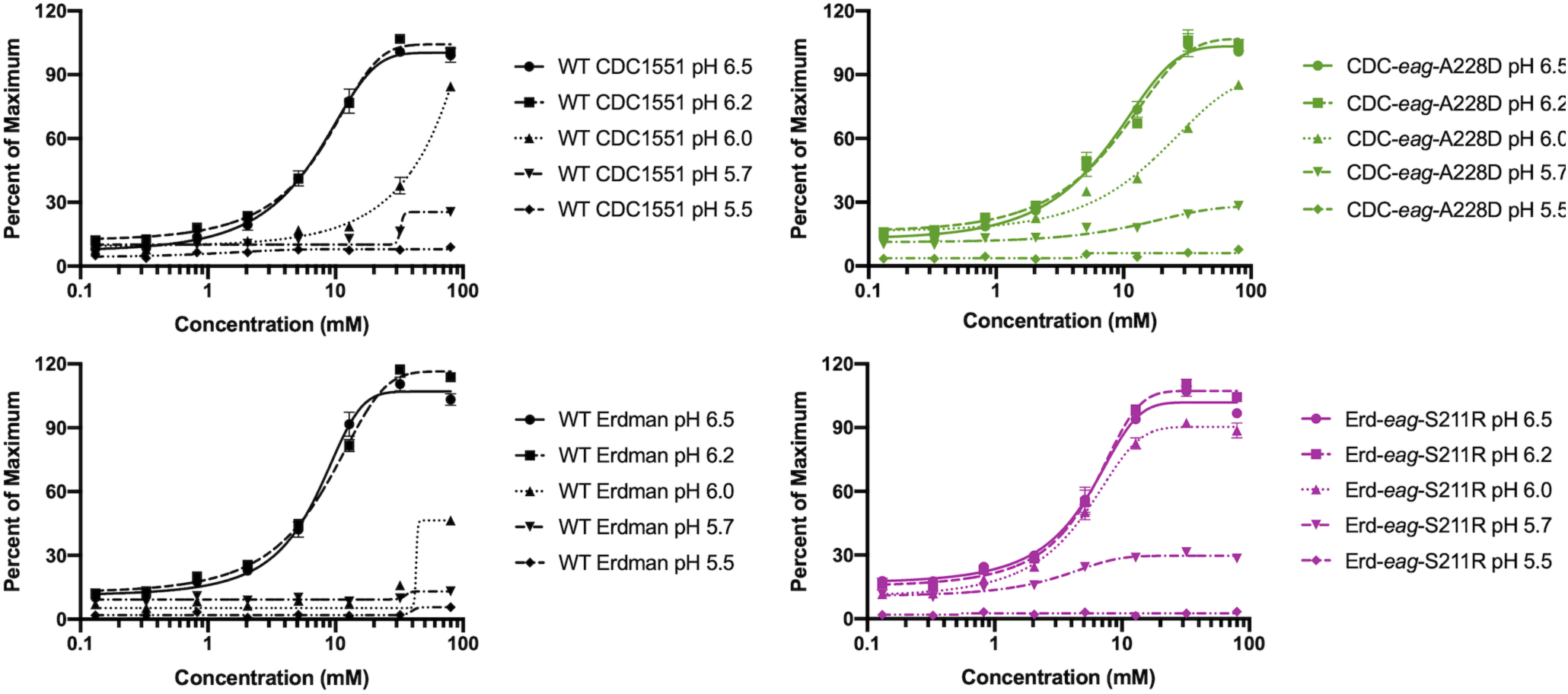
Mtb growth restriction and rescue at low pH is also observed in the native eag variants in CDC1551 and Erdman. WT CDC1551 and Erdman exhibit a slight rescuing of growth at high glycerol concentrations (∼80 mM) in minimal media buffered to pH 5.7, consistent with what has been observed in the Δppe51 complemented strains. The native CDC1551 eag variant, A228D, and native Erdman eag variant exhibit a greater capacity for growth at pH 5.7 at lower concentrations of glycerol (∼5mM) compared to their respective WT strains. Growth analyses were performed at Day 14 following initial inoculation with data being shown as percent of the maximum well-growth. All conditions were conducted in duplicate and representative of multiple independent experiments.

**Supplemental Figure 12.**
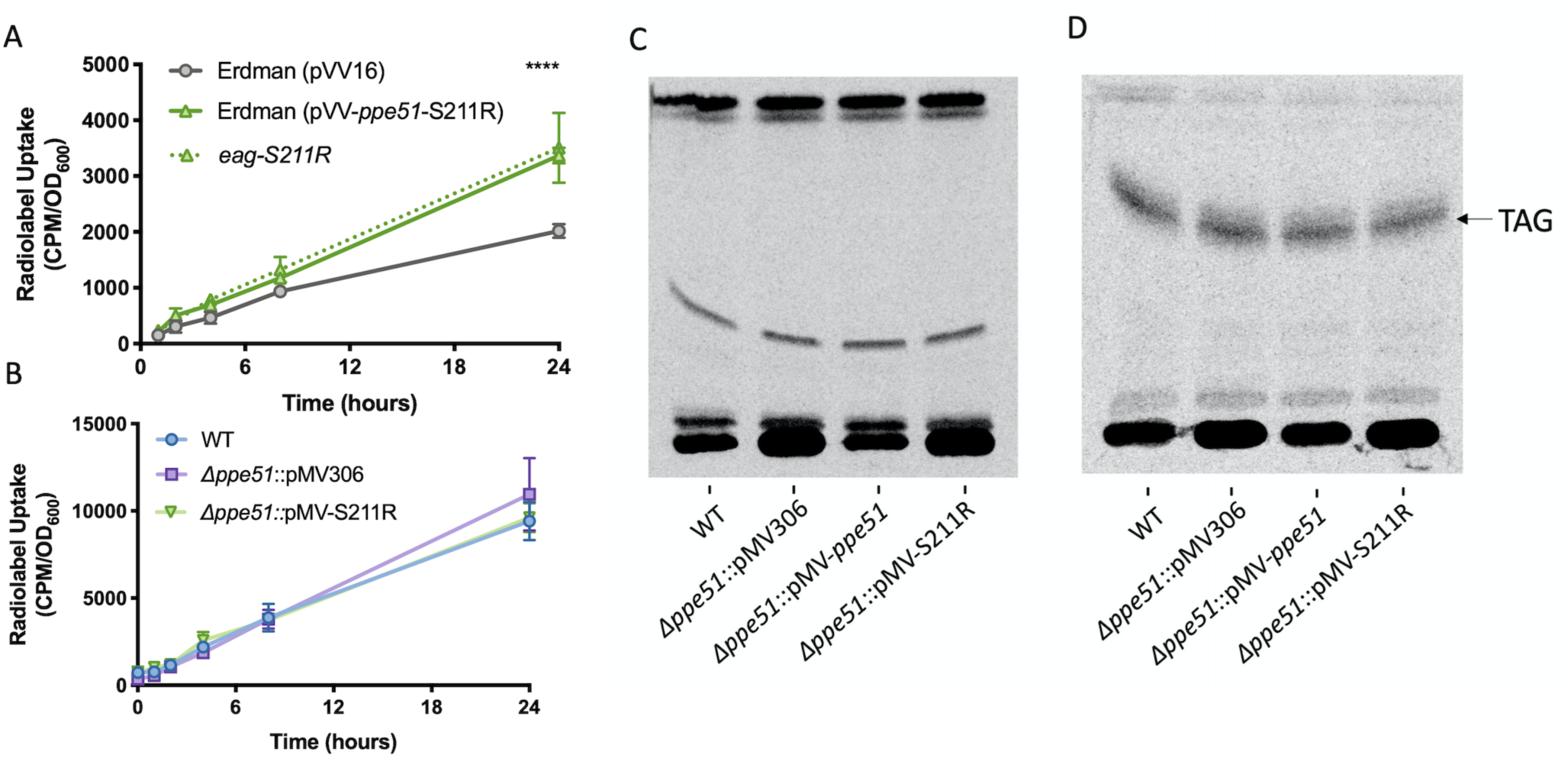
Glycerol uptake in native eag variants (pH 5.7), and radiolabeled uptake and incorporation into lipids at pH 7.0. A) The Erdman native eag S211R variant and the overexpressing pVV16-S211R variant uptake 14C-glycerol at a similar enhanced rate compared to WT Erdman which exhibits a more reduced capacity for radiolabeled uptake. Mtb was pre-adapted for 3 days in minimal media (pH 5.7) with 10 mM glycerol and subsequently washed prior to the additional of radiolabeled glycerol. 14C-glycerol uptake was measured at various timepoints using Scintillation counting over 24 hours. Significance was determined by two-way ANOVA (Tukey’s multiple comparisons test; ****P < 0.0001) B) Erdman WT, Δppe51 (empty vector) and, Δppe51::pMV-S211R were assessed for 14C-glycerol uptake in minimal media buffered to pH 7.0. Strains were pre-adapted for 3 days in minimal media buffered to pH 7.0 with 10 mM glycerol. All strains showed similar rates of glycerol uptake. C) Incorporation of 14C-glycerol into sulfolipids at neutral pH. Sulfolipid is absent at pH 7.0 which is consistent with previous observations made by Baker et. al. Strains were analyzed in duplicate with representative results being shown. D) Incorporation of 14C-glycerol into TAG at acidic pH. TAG is indicated with an arrow and are present in all strains. Strains were analyzed in duplicate with representative results being shown.

**Supplemental Figure 13.**
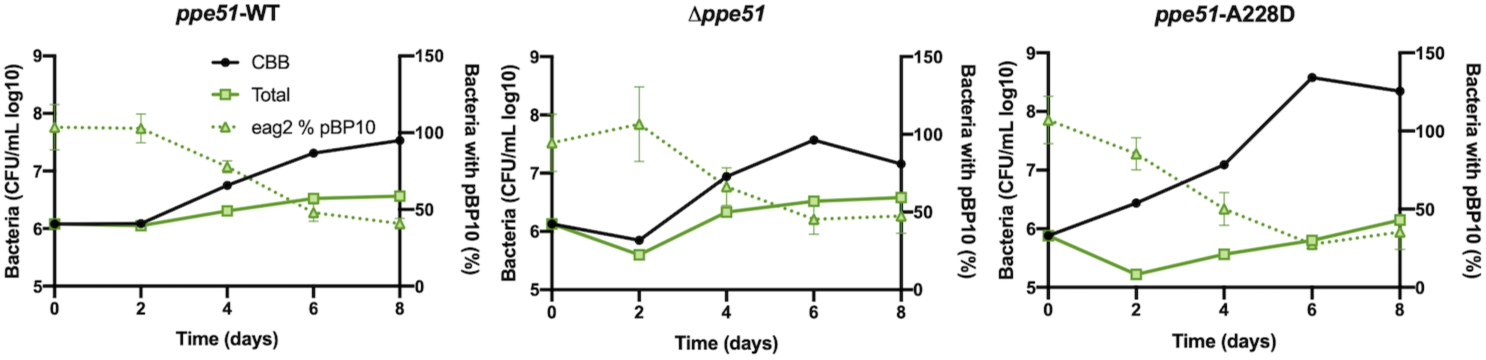
Resting BMDMs infected with native WT CDC1551, Δppe51, and A228D variant strains containing the pBP10 replication clock plasmid. CFUs on selective plates were compared to CFUs on nonselective plates and used to calculated frequency of plasmid-bearing bacteria (% pBP10), cumulative bacterial burden (CBB) of total live and dead bacteria, and total enumerated colonies on nonselective plates (CFU).

**Supplemental Figure 14.**
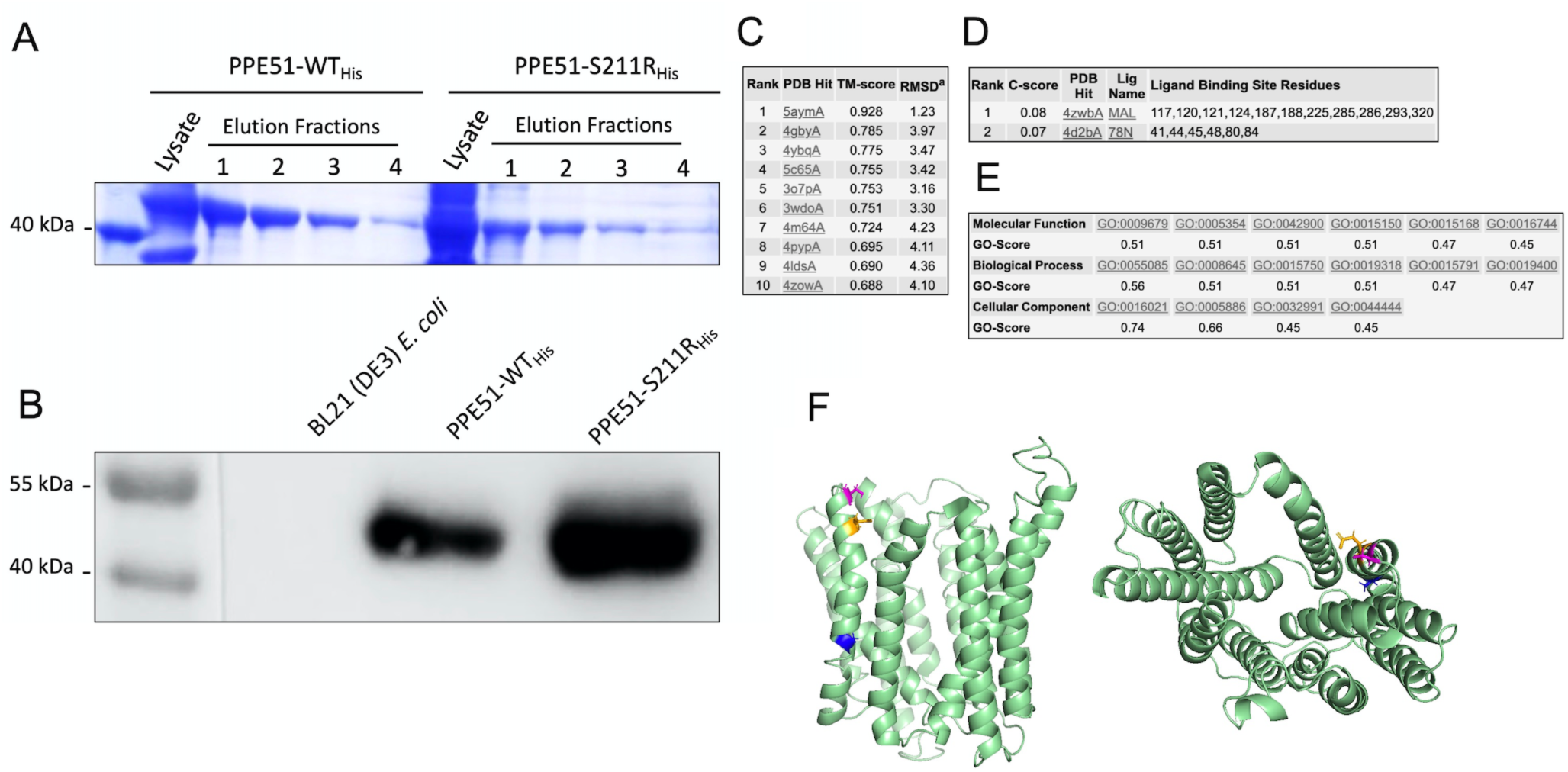
Protein expression of PPE51His and in silico modeling. Lysates of overexpressed His-tagged PPE51-WT and PPE51-S211R were run on a Talon resin column, and fractions were collected in (4) 1 mL aliquots. PPE51-WT and PPE51-S211R proteins were separated on 12% SDS-PAGE gels, which were either stained with Coomassie Blue dye (A) or used for western blots (B). Western blots were incubated with mouse anti-His tag monoclonal antibody followed by HRP-conjugated anti-mouse IgG secondary antibody. The molecular weights of the protein standards are shown on the left. C) Top ten PDB structures close to the target protein. TM-scores are a measurement of the structural similarity between the query structure and known structures in the PDB library in the range [0,1]. TM-scores >0.5 indicate a more correct topology. RMSD is the measurement of the average distance of residues between two structures. D) Ligand binding site prediction. C-score is the confidence score of the prediction in the range [0,1]. A higher score indicates a more reliable prediction. Ligand names are possible binding ligands foud in the BioLiP database. E) Gene ontology (GO) term prediction. Summary prediction of the most common GO terms occurring in three functional aspects (molecular function, biological process, and cellular component). GO-Score is a confidence score of the predicted GO term. A GO-Score >0.5 indicates a more reliable prediction. F) Modeling of the eag mutation sites in PPE51 shows that S211R (pink), E215K (orange), and A228D (blue) all occur on on the same alpha helix.

**Supplemental Figure 15.**
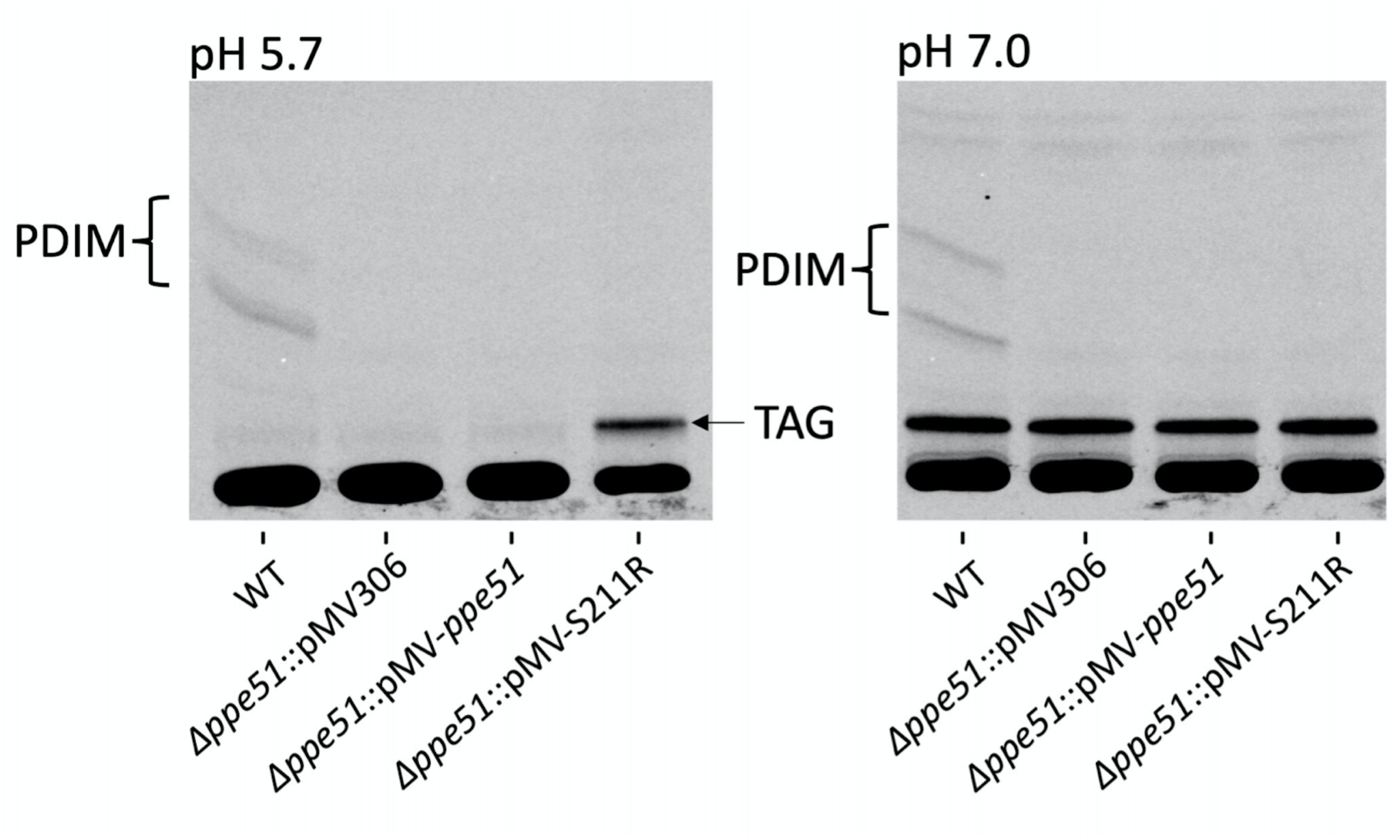
Incorporation of 14C-glycerol into PDIM at acidic and neutral pH. PDIM is indicated with a bracket and accumulates in the WT strain at both pH 5.7 and pH 7.0 but is absent in the ppe51 knockout mutant. A band consistent with TAG appears in Δppe51::pMV-S211R and is indicated by an arrow. Strains were analyzed in duplicate with representative results being shown.

**Table S1.** Mass Spectrometry results for bands associated with PPE51 induction.

| Sample Name | Cluster | Protein Name | Protein accession numbers | Protein molecular weight (Da) | Protein identification probability | Exclusive unique peptide count | Exclusive unique spectrum count | Total spectrum count |
| --- | --- | --- | --- | --- | --- | --- | --- | --- |
| Induced | Cluster of PPE family protein [Mycobacterium tuberculosis CDC1551] (AAK47561.1) | PPE family protein [Mycobacterium tuberculosis CDC1551] | AAK47561.1,KBN10067.1,WP_003416381.1,sp\|P9WHY2.1\|PPE51_MYCTO | 37,979.90 | 100.00% | 5 | 13 | 32 |
| Pellet | Cluster of PPE family protein [Mycobacterium tuberculosis CDC1551] (AAK47561.1) | PPE family protein [Mycobacterium tuberculosis CDC1551] | AAK47561.1,KBN10067.1,WP_003416381.1,sp\|P9WHY2.1\|PPE51_MYCTO | 37,979.90 | 100.00% | 7 | 17 | 54 |

**Table S2.**
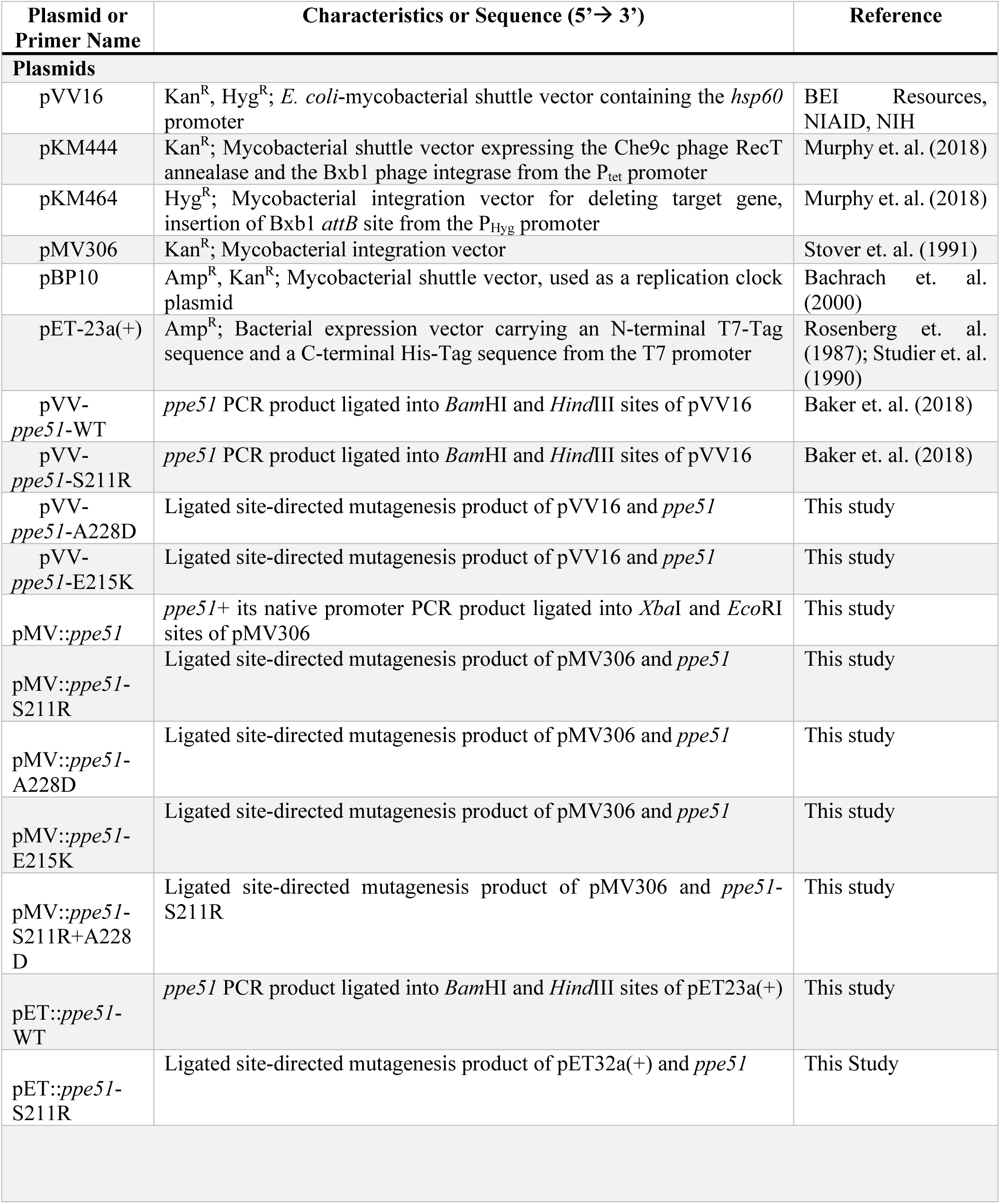
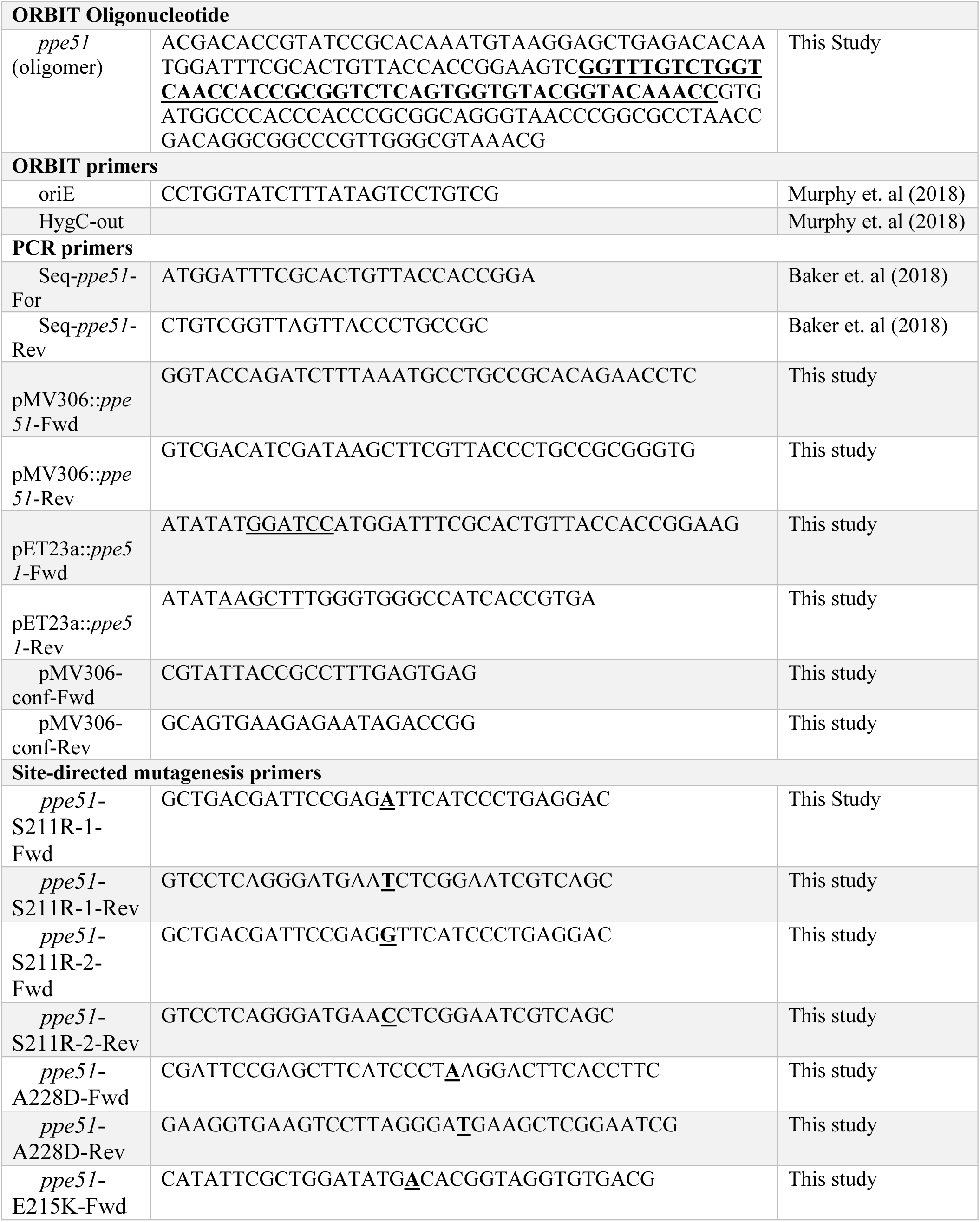
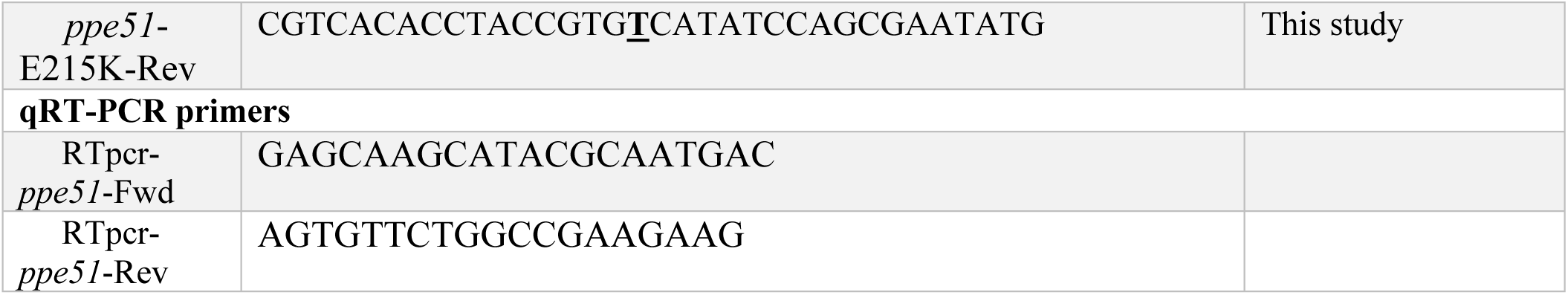
Plasmids and Primers used in this study. The Bxb1 phage *attP* sequence is in bold. Enzyme restriction cut sites are underlined. Site-directed mutagenesis sites are bold and underlined.

